# Redefining the roles of UDP-glycosyltransferases in auxin metabolism and homeostasis during plant development

**DOI:** 10.1101/2021.01.26.427012

**Authors:** Eduardo Mateo-Bonmatí, Rubén Casanova-Sáez, Jan Šimura, Karin Ljung

**Author notes:** Corresponding authors: Eduardo Mateo-Bonmatí and Karin Ljung. These authors equally contributed to this work (Eduardo Mateo-Bonmatí, Rubén Casanova-Sáez). John Innes Centre, NR4 7UA, Norwich, United Kingdom (Eduardo Mateo-Bonmatí).

## Abstract

The levels of the important plant growth regulator indole-3-acetic acid (IAA) are tightly controlled within plant tissues to spatiotemporally orchestrate concentration gradients that drive plant growth and development. Metabolic inactivation of bioactive IAA is known to participate in the modulation of IAA maxima and minima. IAA can be irreversibly inactivated by oxidation and conjugation to Aspartate and Glutamate. Usually overlooked because its reversible nature, the most abundant inactive IAA form is the IAA-glucose (IAA-glc) conjugate. Glycosylation of IAA is reported to be carried out by the UDP-glycosyltransferase 84B1 (UGT84B1), while UGT74D1 has been implicated in the glycosylation of the irreversibly formed IAA catabolite oxIAA. Here we demonstrate that both UGT84B1 and UGT74D1 modulate IAA levels throughout plant development by dual IAA and oxIAA glycosylation. Moreover, we identify a novel UGT subfamily whose members modulate IAA homeostasis during skotomorphogenesis by redundantly mediating the glycosylation of oxIAA.

## INTRODUCTION

Indole-3-acetic acid (IAA), the major natural auxin in plants, coordinates developmental programs throughout the plant's life cycle by integrating external and internal signals into regulated plant responses (Zažímalová *et al.*, 2014; Casanova-Sáez *et al.*, 2021). This is achieved by the formation of concentration gradients that establish auxin maxima and minima within plant tissues. In combination with inter- and intracellular transport, IAA metabolism modulates auxin gradients and is therefore critical for plant growth (Casanova-Sáez *et al.*, 2021). In addition to the many layers of regulation of IAA biosynthesis, redundant mechanisms exist to keep about 75% of the pool of IAA molecules as either transient storage forms or catabolites, thus providing a robust and rapid-response system to fine-tune IAA levels (Ludwig-Muller, 2011). IAA is inactivated in Arabidopsis mainly by irreversible oxidation to oxIAA (2-oxoindole-3-acetic acid) facilitated by DIOXYGENASE FOR AUXIN OXIDATION 1 (DAO1) and DAO2 (Porco *et al.*, 2016), while reversible conjugates can be formed via amide and ester bonds with amino acids and methyl groups by the action of GRETCHEN HAGEN 3 (GH3) and IAA carboxyl methyltransferase (IAMT) respectively (Qin *et al.*, 2005; Staswick *et al.*, 2005). The most abundant reversible IAA inactive forms, however, are IAA conjugates with sugars such as glucose, as observed by direct quantification of IAA metabolites in different plant species (Porco *et al.*, 2016; Pěnčík *et al.*, 2018; Brunoni *et al.*, 2020)

Sugar conjugation confers higher stability and water solubility and has been considered to be a biological tagging mechanism controlling metabolite activity and compartmentalization (Jones & Vogt, 2001). UDP-glycosyltransferases (UGTs) catalyse the transfer of uridine-diphosphate-activated monosaccharides to a variety of compounds, including anthocyanins (Yonekura-Sakakibara *et al.*, 2012), cell wall components (Lin *et al.*, 2016), fatty acids (Rocha *et al.*, 2016), flavonoids (Li *et al.*, 2018), glucosinolates (Grubb *et al.*, 2014) and phenylpropanoids (Sinlapadech *et al.*, 2007). In Arabidopsis, UGTs comprise a gene superfamily of 115 members (Yu *et al.*, 2017), clustered in 19 families (71-91) and 7 subfamilies (A-F), with up to 11 members per subfamily. Several UGTs have also been found to modulate the metabolism of different phytohormones by glycosylation of the bioactive form: UGT71B6 for abscisic acid (Priest *et al.*, 2006); UGT73C5 and UGT73C6 for brassinosteroids (Poppenberger *et al.*, 2005; Husar *et al.*, 2011); UGT85A1, UGT76C1 and UGT76C2 for cytokinins (Smehilova *et al.*, 2016); UGT76E1 for jasmonic acid (Haroth *et al.*, 2019); and UGT89A2 and UGT76D1 for salicylic acid (Li *et al.*, 2014; Chen & Li, 2017; Huang *et al.*, 2018).

A group of UGTs have been suggested as playing a role in the reversible conversion of IAA to IAA-glucose (IAA-glc). Recombinant UGT84B1, UGT84B2, UGT75B1, UGT75B2, and UGT74D1 were found to be able to glycosylate IAA and other natural and synthetic auxins such as IPA (indole-3-propionic acid), IBA (indole-3-butyric acid), and NAA (naphthalene acetic acid) (Jackson *et al.*, 2001; Jin *et al.*, 2013) *in vitro*. UGT74D1 was further found to glycosylate oxIAA *in vitro* and it was suggested that it performs this function *in vivo* (Tanaka *et al.*, 2014). However, the strongest IAA glycosyltransferase activity was observed with UGT84B1 (Jackson *et al.*, 2001). Besides *in vitro* evidence, plants overexpressing *UGT84B1* showed not only higher levels of IAA and IAA-glc, but also a wrinkled and curling leaf phenotype similar to those of other auxin-accumulating mutants (Jackson *et al.*, 2002).

Despite the biochemical evidence supporting a role for UGT84B1 in IAA glycosylation, it was recently suggested that UGT84B1 may also glycosylate oxIAA, on the basis of *in bacteria* assays (Brunoni *et al.*, 2019). Here, we generate a CRISPR/Cas9-based knock-out allele of *UGT84B1* and show, by tissue-specific auxin metabolite profiling and feeding experiments with isotope-labelled [^13^C_6_]IAA, that both UGT84B1 and UGT74D1 modulate IAA levels throughout the plant lifecycle by playing a dual role in IAA and oxIAA glycosylation. We additionally identify a new subfamily of UGTs that have a role in IAA homeostasis during skotomorphogenesis.

## RESULTS

### IAA-glucose accumulates in root tissues while *UGT84B1* is mostly expressed during seed development

IAA-glc has been detected in many plant species, at particularly high abundance in seeds, and it is believed to constitute the main source of IAA during seedling germination and establishment (Ljung *et al.*, 2001). However, the role of this conjugate at later developmental stages is unclear. To further understand the post-germination role of IAA-glc, we revisited datasets of tissue-specific quantification of IAA-glc levels in Arabidopsis (Porco *et al.*, 2016), and observed that roots accumulate IAA-glc at more than 3-fold higher levels compared to other seedling tissues such as hypocotyls, cotyledons, first leaves and shoot apex (Figure 1a). Because the UDP-glycosyltransferase 84B1 (UGT84B1) is believed to account for IAA-glc formation in the plant (Jackson *et al.*, 2001; Jackson *et al.*, 2002), we explored whether the expression pattern of *UGT84B1* parallels the abundance of IAA-glc in plant tissues. We retrieved expression profiling data for *UGT84B1* from Genevestigator (Hruz *et al.*, 2008) using only datasets corresponding to wild-type tissues. According to these data, transcripts of *UGT84B1* are present at very low levels during the vegetative phase, while they peak specifically in some reproductive structures such as endosperm, seeds and siliques (Figure 1b). To validate these data under our experimental conditions, we analysed the expression of *UGT84B1* in young seedlings and siliques by qRT-PCR. Transcript levels of *UGT84B1* were found to be more than 30-fold higher in siliques than in seedlings (Figure 1c), thus confirming the published findings.

**Figure 1.**
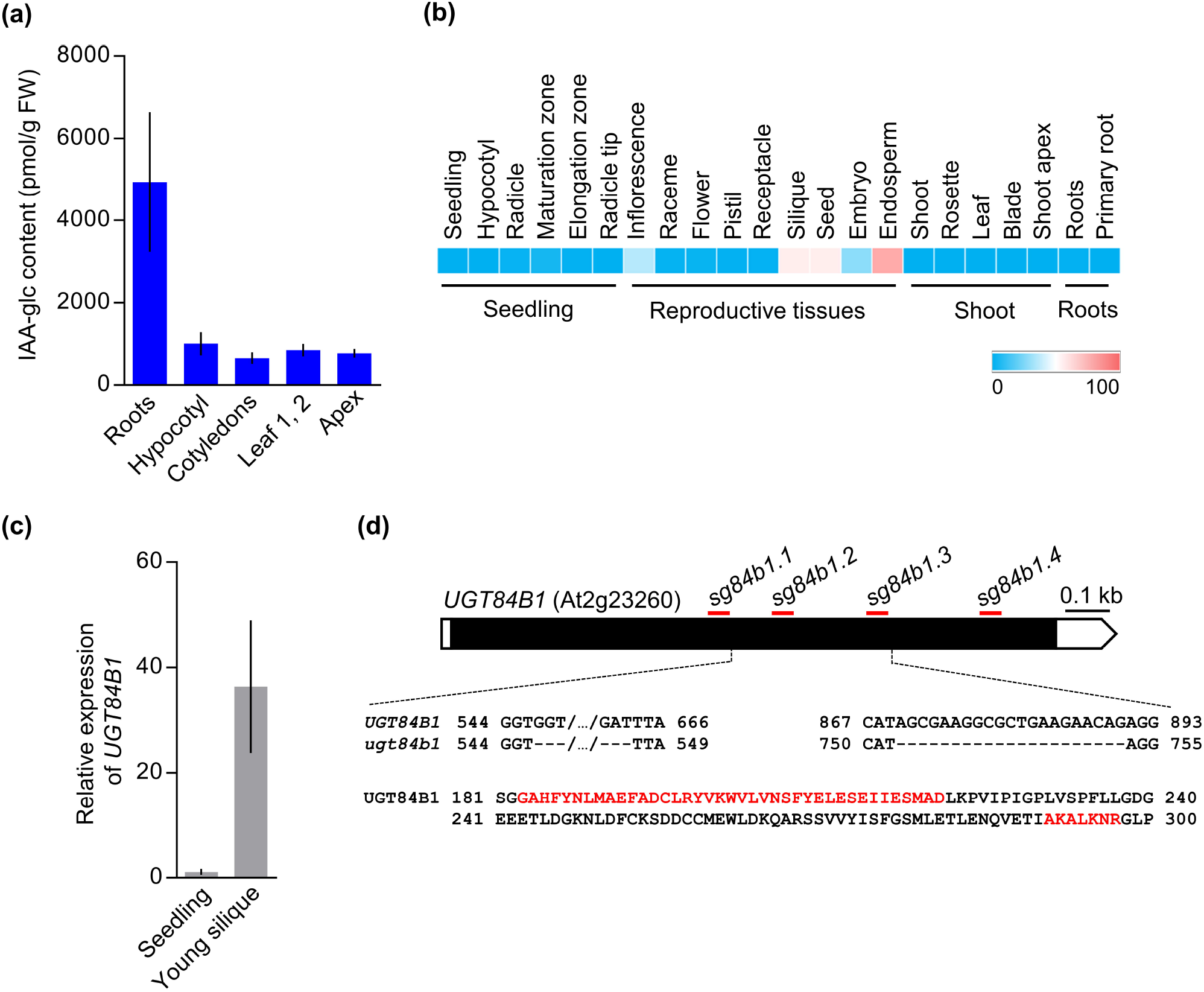
IAA-glc contents do not correlate with *UGT84B1* expression in vegetative tissues. (a) IAA-glucose levels in different tissues of Col-0 plants. Data retrieved from Porco *et al.* 2016. (b) Expression pattern of the *UGT84B1* gene in different tissues and organs. Transcriptomic data were obtained from Genevestigator datasets corresponding to wild-type tissues. Color scale indicates percentage of expression potential. (c) Relative expression of the *UGT84B1* gene in wild-type seedlings and young siliques. Bars indicate relative expression of *UGT84B1* in 7-day-old Col-0 seedlings and in young siliques from 39-day-old Col-0 plants. Error bars indicate the interval delimited by 2^−^ΔΔCT±SD. (d) Structure of the *UGT84B1* gene. Open and black boxes represent untranslated and translated regions respectively. Red horizontal lines represent the positions of the nucleotide sequences (not drawn to scale) used to design the guide RNAs for CRISPR/Cas9-based gene editing. *ugt84b1* plants carry two nucleotide deletions (positions 547-663 and 870-890) that generate two protein deletions (positions 183-221 and 291-297; highlighted in red). Scale bar indicates 100 bp.

### Tissue-specific auxin metabolite profiling of *ugt84b1* mutants uncovers a role for UGT84B1 in oxIAA glycosylation throughout plant development

To further understand the role of UGT84B1 in IAA metabolism during the Arabidopsis lifecycle, and because no loss-of-function allele was available at the time this study was initiated, we generated a CRISPR/Cas9-based *ugt84b1* knockout mutant carrying two deletions in the protein sequence (amino acids 183-221 and 291-297; Figure 1d). In line with other recently reported *UGT84B1* knockout alleles (Aoi *et al.*, 2020), *ugt84b1* mutant plants did not show any obvious developmental defect (Supplementary Figure 1).

To determine whether UGT84B1 contributes to IAA homeostasis *in planta*, we quantified the levels of IAA and oxIAA, their glycosylated forms IAA-glc and oxIAA-glc, and the irreversibly synthesized catabolites IAA-Asp and IAA-Glu in *ugt84b1* plants. Given the differences in *UGT84B1* expression and IAA-glc contents across plant tissues (Figure 1a), we decided to analyse three different tissues: roots and shoots from 7-days-old seedlings and young siliques from 39-day-old plants (Figure 2). In line with the reported role of UGT84B1 in IAA glycosylation (Jackson *et al.*, 2001; Jackson *et al.*, 2002; Aoi *et al.*, 2020) and the specific expression of the gene in siliques (Figure 1c), we found decreased levels of IAA-glc in *ugt84b1* siliques compared to Col-0 (Figure 2), while IAA-glc concentrations were higher in young roots and shoots of *ugt84b1*. Strikingly, the levels of oxIAA-glc were decreased in the *ugt84b1* mutant in all tissues analysed (Figure 2). IAA homeostasis was significantly affected by the loss of *UGT84B1* in all tissues, as indicated by the decreased IAA contents of the *ugt84b1* mutant tissues (Figure 2) and by the altered contents of the catabolites oxIAA, IAA-Asp and IAA-Glu, notably in siliques (Supplemental Figure 2). Overall, these results suggest that UGT84B1 is additionally required for oxIAA glycosylation and auxin homeostasis during plant development, and strongly favours the hypothesis that other UGT members may function redundantly in IAA glycosylation at the seedling stage.

**Figure 2.**
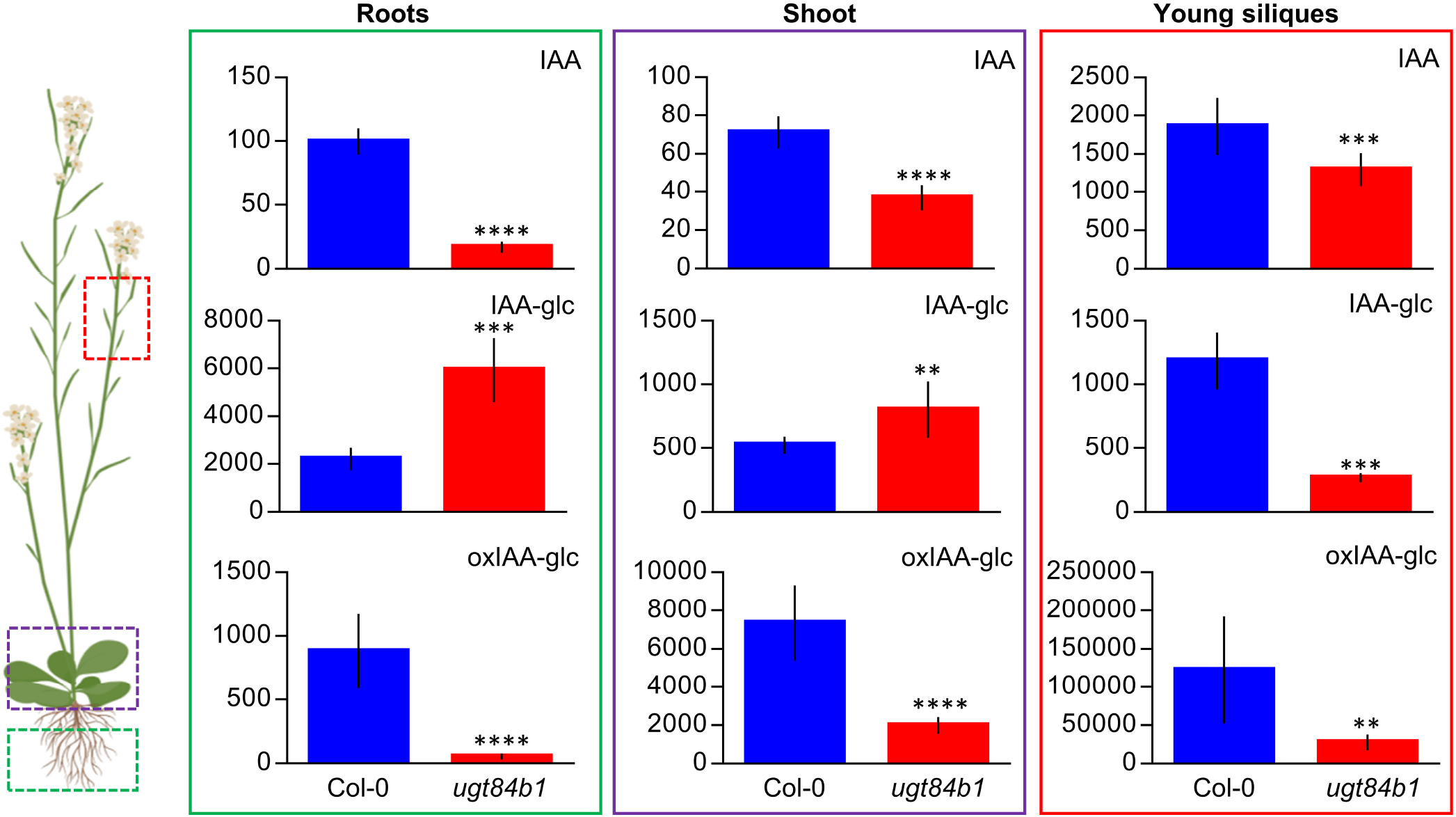
Tissue-specific profiling of IAA metabolites in the *ugt84b1* mutant. IAA and the IAA metabolites IAA-glucose (IAA-glc) and oxIAA-glucose (oxIAA-glc) were quantified in roots (green outline) and shoots (purple outline) from 7-day-old seedlings, and in young siliques (red outline) from 39-day-old plants. The concentrations of all metabolites are given in picomoles per gram of fresh weight. Samples were analysed with ten independent biological replicates, and error bars represent the standard deviation (SD). Asterisks indicate statistically significant differences from Col-0 (\*\**p* < 0.01, *** *p* < 0.001, and **** *p* < 0.0001; Student’s *t* test).

### Searching for new UGTs involved in IAA glycosylation

In order to find additional members of the UGT superfamily with functions in IAA glycosylation and homeostasis, we performed an *in silico* search for IAA-inducible UGTs. We employed the dataset described in (Lewis *et al.*, 2013) in which the authors exposed wild-type roots to 1 μM IAA, and retrieved for analysis those UGTs whose function had not yet been established (Figure 3a). Among all these UGTs, only *UGT76E5* showed constant and strong induction upon IAA treatment, while *UGT86A2* or *UGT90A2* showed lesser induction (Figure 3a). We therefore focused on *UGT76E5* for further experiments. To validate the microarray data under our experimental conditions, we used qRT-PCR to analyse the expression of *UGT76E5* in 7-day-old seedlings treated with 1 μM of IAA. The relative expression of *UGT76E5* after 4 hours of treatment was around 5-fold higher than in mock-treated plants (Figure 3b). Because *UGT76E5* is a non-characterized member of this superfamily, we used the ATTEDII tool to analyse the *UGT76E5* co-expression network. Based on array datasets, we found a gene involved in IAA inactivation, *GH3.1* (At2g14960), within the co-expression map (Supplementary Figure 3a). Additional co-expression analyses based on RNA-seq datasets indicated that *GH3.17* (At1g28130), *SMALL AUXIN UP-REGULATED RNA*-like 55 (*SAUR-like 55*; At5g50760), and *SAUR21* (At5g01830) are also co-expressed with *UGT76E5*, thus supporting a relationship between *UGT76E5* and IAA homeostasis (Supplementary Figure 3b).

**Figure 3.**
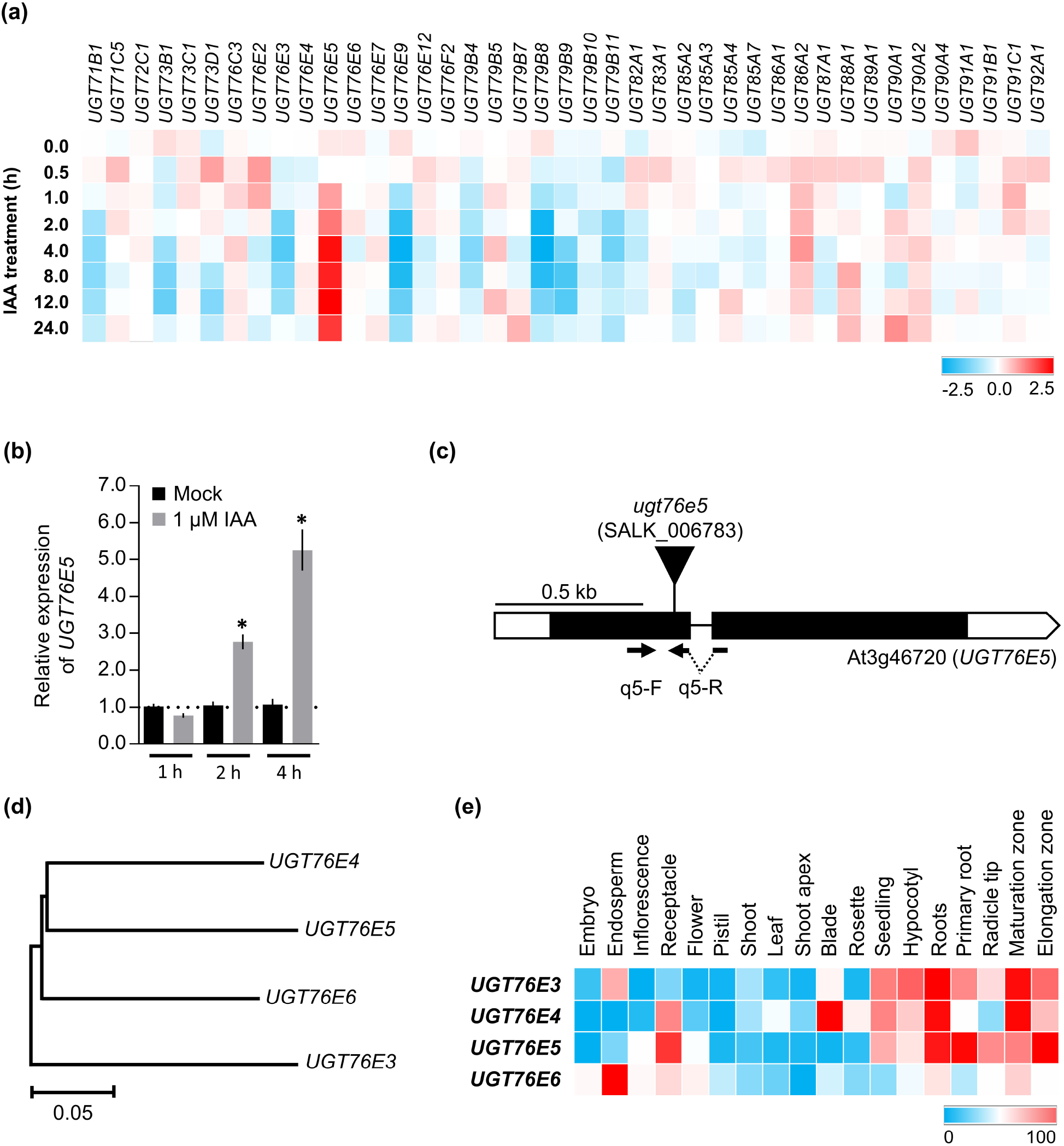
A non-characterized UGT family member is upregulated by auxin in roots. (a) Transcriptional responses of all uncharacterized Arabidopsis UGT genes to 1 μM IAA in root tissues. Transcriptomic data were obtained from the GSE42007 dataset at Genevestigator. (b) Relative expression analysis of *UGT76E5* in response to 1 μM IAA treatments. Bars indicate relative expression of *UGT76E5* in 7-day-old Col-0 seedlings after mock (black) and IAA (grey) treatments at different times. Error bars indicate the interval delimited by 2^−^ΔΔCT±SE. Asterisks indicate ΔCT values significantly different from those of the mock treatment in a Mann-Whitney *U* test (\**p*<0.001; n=9). (c) Structure of the *UGT76E5* gene indicating the position of the *ugt76e5* mutation. Boxes and lines represent exons and introns respectively. Open and black boxes represent untranslated and translated regions respectively. The triangle indicates a T-DNA insertion. Horizontal arrows represent the oligonucleotides q_UGT76E5_F (q5-F) and q_UGT76E5_R (q5-R), not drawn to scale, used as primers in (b). (d) Neighbour-joining tree of proteins of the UGT76E subfamily. The tree was constructed with MEGAX (see methods) and inferred from 1000 replicates. (e) Expression patterns of the *UGT76E3*, *UGT76E4*. *UGT76E5*, and *UGT76E6* genes in different tissues. Transcriptomic data were obtained from Genevestigator datasets corresponding to wild-type tissues. Scale bars indicate (c) 0.5 kilobases and (d) 5% amino acid sequence changes according to the Poisson correction method (Nei and Kumar, 2000). Genevestigator colour scales indicate (a) log2-ratio values from −2.5 to 2.5 and (e) percentage of expression potential.

In contrast to *UGT84B1*, *UGT76E5* expression was detected in different vegetative organs, primarily in roots and hypocotyls (Figure 3e). To study the role of *UGT76E5* in IAA metabolism, we obtained a T-DNA allele harbouring an insertion (SALK_006783) in the first exon of *UGT76E5,* which we named *ugt76e5* (Figure 3c). We also generated transgenic plants expressing the *UGT76E5* transcriptional unit under the control of a double 35S promoter (seven independent families with high *UGT76E5* expression level were obtained; Supplementary Figure 4). Neither *ugt76e5* nor *35S_pro_:UGT76E5* homozygous plants showed any noticeable morphological phenotype (Supplementary Figure 1). BLASTP searches allowed us to identify three close paralogs of *UGT76E5* in the Arabidopsis genome, *UGT76E3* (At3g46700), *UGT76E4* (At3g46690), and *UGT76E6* (At3g46680) (Figure 3d), whose redundancy might explain the lack of morphological defects in *ugt76e5* plants. Like *UGT76E5*, these paralogs are expressed in vegetative organs and root tissues, with the exception of *UGT76E6* whose expression is reminiscent of that of *UGT84B1* and is largely restricted to the endosperm (Figure 3e). Phylogenetic analyses indicate that these four paralogs (hereafter *UGT76E3456*) fall into the same clade of the UGT76 family (Supplementary Figure 5) and multiple alignment of the UGT76E3456 protein sequences revealed a high degree of amino acid identity among them (Supplementary Figure 6). Finally, we found an IAA amido synthetase, *GH3.17*, and several *SAUR* and *SAUR-like* auxin responsive genes among the co-expression networks of *UGT76E3, UGT76E4, and UGT76E6* (Supplementary Figure 3b). Taken together, the auxin-inducible expression of *UGT76E5* along with the co-expression analyses of the *UGT76E3456* family members suggest involvement of this subfamily in IAA homeostasis.

### Feeding experiments with [^13^C_6_]IAA reveal a dual role for UGT84B1 and UGT74D1 in IAA and oxIAA glycosylation, and a minor contribution by UGT76E3456

We next aimed to explore the role of UGT76E3456 in auxin metabolism in Arabidopsis seedlings. Because the four *UGT76E* genes are clustered together on chromosome 3 (Supplementary Figure 7a), we followed a CRISPR/Cas9-based approach to simultaneously inactivate the four paralogs. We found a genomic sequence with the potential to concurrently target *UGT76E3*, *UGT76E4* and *UGT76E5* (*UGT76E345*) (Supplementary Figure 8a). To generate the quadruple *ugt76e3456* knockout, we transformed the CRISPR/Cas9 construct into *ugt76e6* plants, which carry a T-DNA insertion (SALK_200519) in the first exon of *UGT76E6* (Supplementary Figure 7b). After screening for edited plants, a single nucleotide insertion was found within the target region in *UGT76E3* and *UGT76E4* (Supplementary Figures 9 and 10) and a 66-nt deletion, encompassing part of the target region, in *UGT76E5* (Supplementary Figure 11). These changes are predicted to truncate both UGT76E3 and UGT76E4 proteins and to generate a 22-amino acid deletion in UGT76E5 (Supplementary Figure 8b). We did not observe any noticeable morphological defect in plants from the quadruple *ugt76e3456* mutant under standard growth conditions (Supplementary Figure 1).

Besides UGT84B1, only one UGT family member has been previously reported to catalyse oxIAA-glc formation *in vivo*, UGT74D1 (Tanaka *et al.*, 2014). According to Genevestigator data, the expression pattern of *UGT74D1* is opposite to that of *UGT84B1*, with expression being higher during vegetative stages, especially in roots, and lower in reproductive tissues (Supplementary Figure 12a). We obtained a T-DNA allele harbouring an insertion (SALK_004870) in the second exon of *UGT74D1*, which generates no wild-type transcript and we named *ugt74d1* (Supplementary Figure 12b-c; Tanaka *et al.*, 2014). Like *ugt84b1* and *ugt76e3456*, these plants did not show any morphological defect but a marginally early flowering (Supplementary Figure 1). To better understand the roles of UGT84B1, UGT74D1, and UGT76E3456 we quantified the levels of IAA and oxIAA, their glycosylated forms IAA-glc and oxIAA-glc, and the catabolites IAA-Asp and IAA-Glu in 7-day-old seedlings of the wild-type Col-0, single mutants *ugt84b1* and *ugt74d1*, the quadruple mutant *ugt76e3456*, and the transgenic *35S_pro_:UGT76E5* seedlings (Supplemental Figure 13). In parallel, we measured the levels of ^13^C_6_-isotope-labelled IAA metabolites in these plants after feeding with [^13^C_6_]IAA (Figure 4).

**Figure 4.**
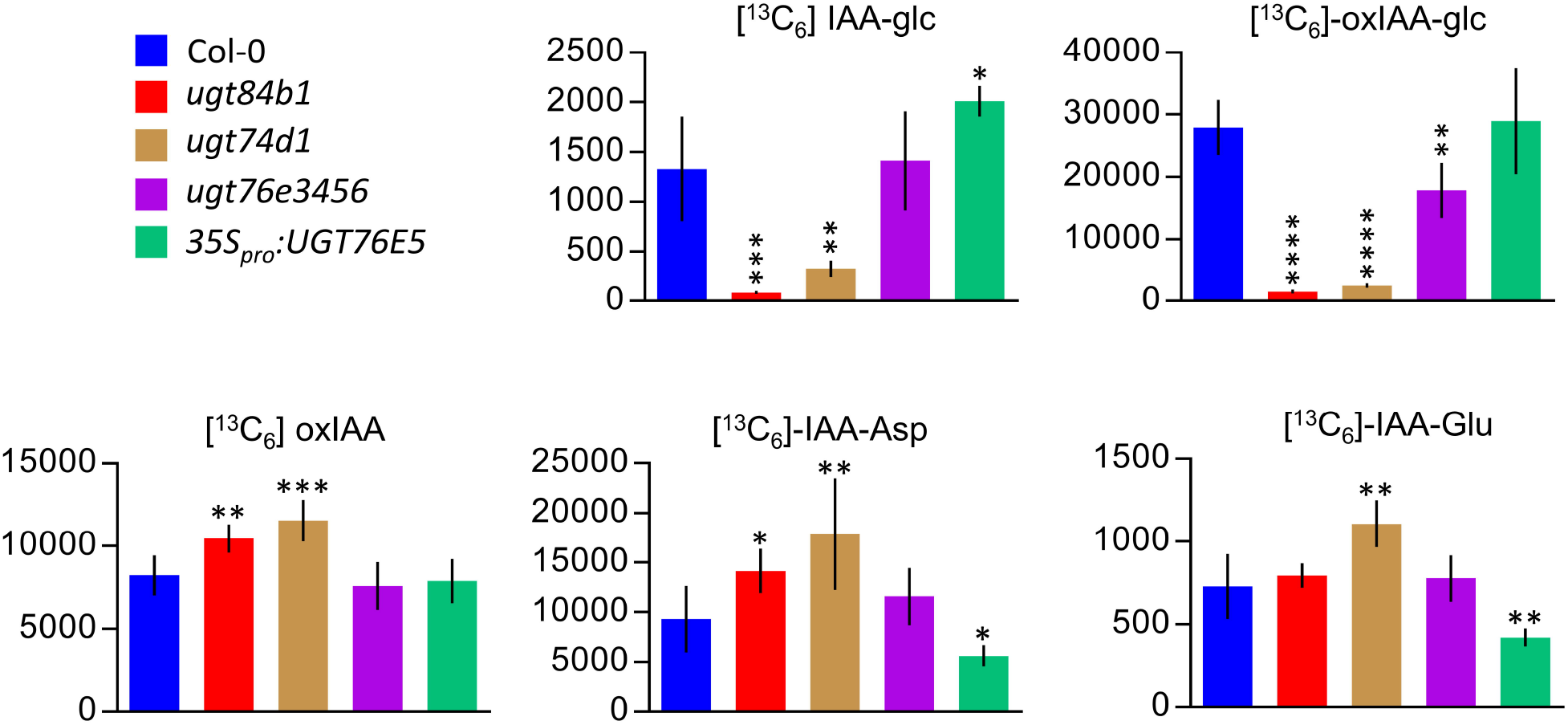
*De novo* synthesis of IAA metabolites in *ugt84b1*, *ugt74d1*, *ugt74d1 ugt84b1*, *ugt76e3456* and *35S_pro_:UGT76E5* plants. Formation of IAA metabolites in 7-day-old seedlings of the genotypes indicated after incubation with [^13^C_6_]IAA for 12 hours. Bars indicate the concentrations (picomoles / g of fresh weight) ± SD of all metabolites. Samples were analysed with five independent biological replicates. Statistically significant differences from Col-0 (\**p* < 0.05, ** *p* < 0.01, \*\*\**p* < 0.001, and \*\*\*\**p* < 0.0001; Student’s *t* test).

In line with our tissue-specific profiling of the *ugt84b1* mutant (Figure 2), lower steady-state levels of IAA and oxIAA-glc, but not IAA-glc, were detected in *ugt84b1* seedlings, again suggesting involvement of UGT84B1 in IAA homeostasis through the regulation of oxIAA-glc formation (Supplemental Figure 13). A similar trend was observed in *ugt74d1* (Supplemental Figure 13). However, the [^13^C_6_]IAA feeding experiment showed that the ability of the *ugt84b1* and *ugt74d1* mutants to synthesize *de novo* both IAA-glc and oxIAA-glc is severely impaired, while oxIAA, IAA-Asp and IAA-Glu formation occurred at a significantly higher rate (Figure 4). These results demonstrate a dual role for UGT84B1 and UGT74D1 in IAA homeostasis through IAA and oxIAA glycosylation *in vivo*.

Slightly lower *de novo* synthesis and steady-state levels of oxIAA-glc were also found in the *ugt76e3456* mutant (Figure 4 and Supplemental Figure 13), while no differences were seen in the *35S_pro_:UGT76E5* transgenic overexpressing line compared to Col-0 at steady state (Supplemental Figure 13). However, *35S_pro_:UGT76E5* plants fed with [^13^C_6_]IAA showed a roughly 2-fold higher IAA-glc formation rate compared to Col-0 and a reduced rate of synthesis of IAA-Asp and IAA-Glu, strongly suggesting that UGT76E5 possesses IAA glycosyltransferase activity *in vivo* and perturbs IAA homeostasis (Figure 4).

### UGT76E3456 control IAA homeostasis during skotomorphogenesis

When kept in darkness, the seedling developmental program is set to a skotomorphogenic mode, in which resources are allocated to hypocotyl elongation in order to find the light (Josse & Halliday, 2008). Several groups have showed that mutations in genes involved in auxin homeostasis alter differential hypocotyl growth (Gray *et al.*, 1998; Sun *et al.*, 2012; Zheng *et al.*, 2016; Chen *et al.*, 2020). To determine whether mutations in any of the UGTs studied in this work affect hypocotyl growth, we measured the hypocotyl length in Col-0, *ugt84b1*, *ugt74d1*, and *ugt76e3456* plants under normal light conditions, shade and darkness. Although the length of light-grown hypocotyls was completely indistinguishable among all genetic backgrounds (Figure 5a), *ugt76e3456* mutant hypocotyls grown in either shade or darkness were found to be significantly longer than those of Col-0 (Figure 5b, 5c).

**Figure 5.**
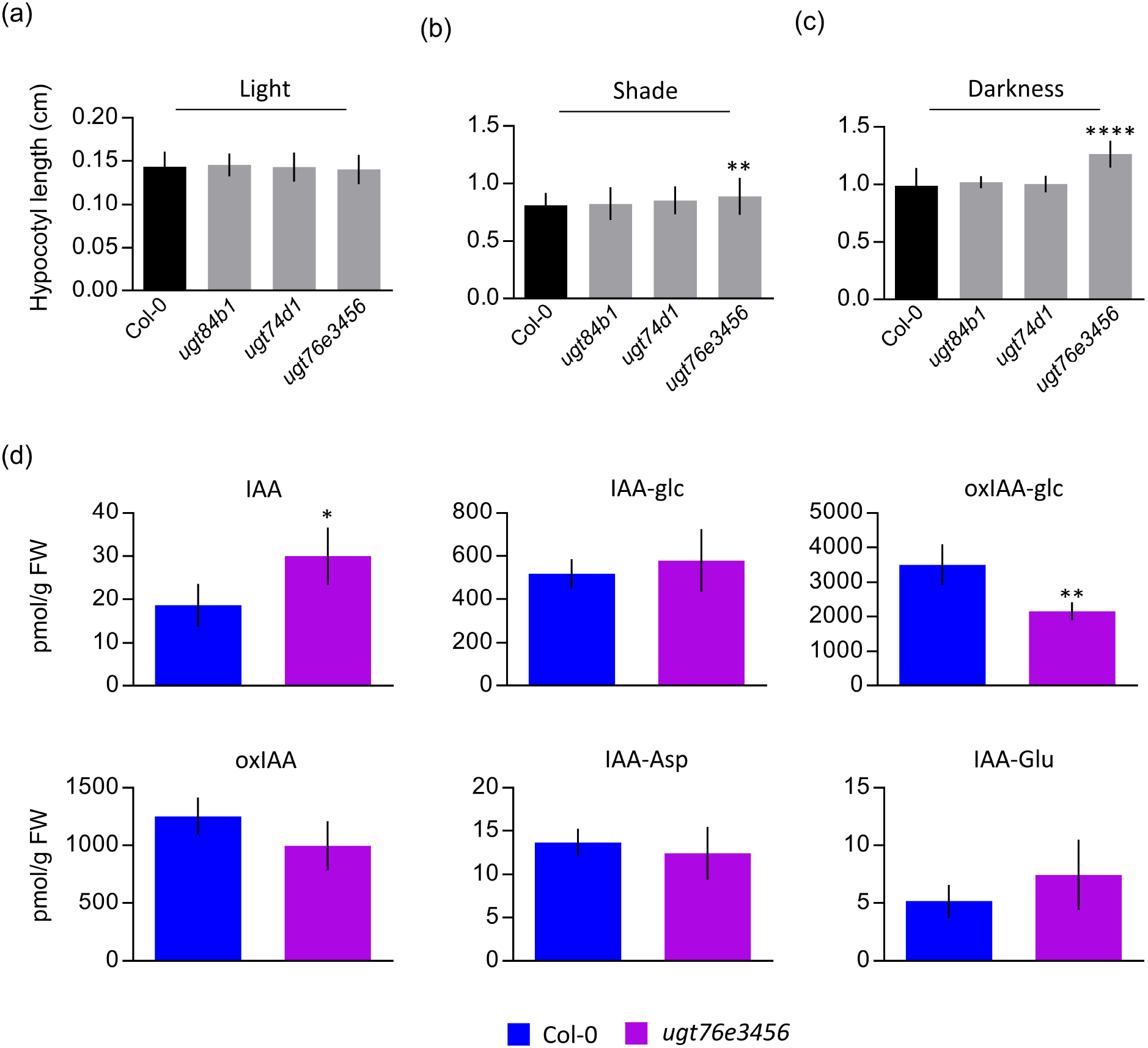
*ugt76e3456* plants show enhanced shade- and darkness-induced hypocotyl elongation. (a-c) Hypocotyl length of Col-0, *ugt84b1*, *ugt74d1*, and *ugt76e3456* seedlings grown in (a) light, (b) shade, and (c) darkness. (d) IAA and IAA metabolite quantification performed in dark-grown hypocotyls. Bars indicate (a-c) length means ± SE (n ≥ 30), (d) concentration (picomoles / g of fresh weight) means ± SD (n = 4). Statistically significant differences from Col-0 (\**p* < 0.05, ** *p* < 0.01, and **** *p* < 0.0001; Student’s *t* test).

To further investigate whether the effect of UGT76E3456 on hypocotyl elongation is connected to modulation of auxin metabolism, we performed auxin metabolite profiling of dark-grown hypocotyls from Col-0 and the *ugt76e3456* quadruple mutant (Figure 5d). In line with their longer hypocotyls, significantly higher levels of IAA were found in the hypocotyls of the *ugt76e3456* quadruple mutant. Consistently with our observations from seedlings fed with labelled IAA (Figure 4), the quadruple mutant showed lower levels of oxIAA-glc in the hypocotyls when grown in darkness (Figure 5d), thus opening up the possibility that these genes play a specific role in oxIAA glycosylation and IAA homeostasis during skotomorphogenesis.

## DISCUSSION

### A redefinition of the roles of UGT84B1 and UGT74D1 in IAA homeostasis

Oxidation and conjugation have been shown to play important roles in the regulation of auxin levels for the control of plant growth and development (Casanova-Sáez *et al.*, 2021). This complex and intertwined net of redundant pathways includes IAA glycosylation, a reversible reaction (through hydrolysis of IAA-glc) that not only participates in temporary inactivation of IAA, but also creates a readily available source of energy and IAA without the need for *de novo* synthesis to trigger fast auxin-mediated responses (Jones & Vogt, 2001).

IAA glycosylation is catalysed by of members of the *UGT* multigene family, which are present from bacteria to humans (Mackenzie *et al.*, 1997). The size of this family has evolved according to organism complexity in the plant kingdom; there are 3 UGTs in the unicellular microalga *Chlamydomonas reinhardtii*, 21 in the moss *Physcomitrella patens*, 115 in Arabidopsis and 200 in *Oryza sativa* (Yu *et al.*, 2017). Despite the challenging size of the family, genetic and biochemical approaches have revealed roles for specific family members in phytohormone glycosylation (Ostrowski & Jakubowska, 2014). *In vitro* screens have uncovered UGT84B1 and UGT74D1 as two potentially major players in IAA glycosylation (Jackson *et al.*, 2001; Jackson *et al.*, 2002; Jin *et al.*, 2013). UGT74D1 was later reported to participate in oxIAA glycosylation in the plant, as a T-DNA *ugt74d1* insertional mutant was found to accumulate oxIAA, while having reduced oxIAA-glc levels (Tanaka *et al.*, 2014). The lack of loss-of-function mutants, however, has hampered a straightforward comparative analysis of the UGT84B1 function *in vivo*. The function of UGT84B1 *in vivo* was reported in overexpression lines nearly 20 years ago (Jackson *et al.*, 2002), but it was not until very recently that the first *ugt84b1* loss-of-function mutant was reported in Arabidopsis and shown to have decreased steady-state levels of IAA-glc in seedlings (Aoi *et al.*, 2020).

Here, we provide the first evidence of a dual role for UGT84B1 in both IAA and oxIAA glycosylation *in vivo*, as demonstrated by the severely impaired *de novo* formation of IAA-glc and oxIAA-glc in plants with our CRISPR/Cas9-based *ugt84b1* knockout allele. Tissue-specific IAA metabolite profiling further revealed a stricter requirement for UGT84B1 for IAA glycosylation in siliques, while the loss of UGT84B1 function appears to be compensated by redundant UGT members in seedling roots and shoots. This agrees with the expression pattern of the *UGT84B1* gene, which is preferentially transcribed in the endosperm. Remarkably, we found the levels of oxIAA-glc to be greatly reduced in seedling roots and shoots, and in siliques, of the *ugt84b1* mutant, which points to UGT84B1 playing a predominant role in oxIAA glycosylation throughout plant development. The profound effects of UGT84B1 on IAA and oxIAA glycosylation in seedling tissues is in remarkable contrast with the relatively low expression of the gene at these stages. While further research will be needed to clarify the causes, that an enzyme mostly transcribed in reproductive tissues has a prevailing role during the vegetative phase represents an exciting challenge to be explored. Because the levels of oxIAA-glc, but not IAA-glc, were previously reported for a *ugt74d1* insertional mutant (Tanaka *et al.*, 2014), we also aimed to explore a potential dual role for UGT74D1 in IAA and oxIAA glycosylation. The metabolic behaviour of *ugt74d1* was very similar to that of *ugt84b1,* with a dramatic impairment of *de novo* synthesis of IAA-glc and oxIAA-glc. This dual role for UGT74D1 was recently suggested by experiments using an heterologous approach in *E. coli* (Brunoni *et al.*, 2019). Our results demonstrate that UGT74D1 modulates IAA levels by IAA and oxIAA glycosylation *in vivo*.

Our tissue-specific metabolite profiling supports the conclusion that both UGT84B1 and UGT74D1 modulate IAA levels during plant development. However, given that glycosylation of IAA and oxIAA involves, respectively, transient and catabolic inactivation of auxin it is surprising that IAA levels are reduced rather than increased in tissues of the *ugt84b1* and *ugt74d1* mutants. It seems plausible, nevertheless, that such regulation of IAA levels reflects a specific compensatory mechanism operating in plants, as overexpression of UGT84B1 has been shown to result in increased IAA levels (Jackson *et al.*, 2002; Aoi *et al.*, 2020). This increase in free IAA levels has also been reported in Arabidopsis plants overexpressing an IAA glycosyltransferase from Maize (Ludwig-Muller *et al.*, 2005). This strongly suggests that the hydrolysable nature of the IAA-glc conjugates, together with the redundant action of additional UGT members, underlie the observed regulation of IAA homeostasis.

### A new set of redundant players locally controlling IAA homeostasis

Large gene families such as the one consisting of UGTs arose from genome duplications (Soltis *et al.*, 2009). After duplication of the ancestral gene, new copies may acquire divergent functions (neofunctionalization); alternatively they can maintain the ancestral function but specialize with respect to the time or the tissue in which they act (subfunctionalization). Considering the short genetic distance between *UGT76E3, UGT76E4, UGT76E5,* and *UGT76E6*, it is likely that they all derive from recent duplication events and act in a redundant manner. Our data indeed suggest that the four paralogs of the UGT76E3456 subfamily play redundant roles in oxIAA glycosylation and IAA homeostasis, which is particularly required for regulating the skotomorphogenic response. Considering the multifunctionality of these enzymes, we cannot completely exclude the possibility that the metabolic and phenotypic effects presented here arise from altering the metabolism of other phytohormones or secondary metabolites. However, our experiments performed both in seedlings and in etiolated hypocotyls consistently showed altered levels of oxIAA-glc in a quadruple *ugt76e3456* mutant. Moreover, the longer dark-grown hypocotyls in the quadruple *ugt76e3456* mutant correlate with decreased oxIAA-glc and increased IAA levels in their hypocotyls. This degree of tissue- and process-specialization has already been reported for GH3.17, a member of the GH3 family of IAA amidosynthetases. Mutations in *GH3.17* block the IAA-Glu pathway, provoking an increment in the IAA content and the length of the hypocotyls (Zheng *et al.*, 2016). More recently, a member of the UGTs (UGT76F1) was found to modulate IAA biosynthesis by glycosylating the major auxin precursor indole-3-pyruvic acid specifically in hypocotyls (Chen *et al.*, 2020). While the connection between the glycosylation of oxIAA, the major IAA catabolite, and the modulation of IAA levels is still unknown, the mutants presented here represent a valuable tool with which to explore this area of research.

It is worth noting that our data on *de novo* formation of isotope-labelled IAA metabolites do not fully exclude a role for *UGT76E5* in IAA glycosylation, as *35S_pro_:UGT76E5* plants were able to produce a 2-fold higher IAA-glc level compared to Col-0. At the same time, the similar steady-state IAA-glc levels found in *ugt84b1*, *ugt74d1* and *ugt76e3456* suggest that other IAA-glycosyltransferases participate redundantly in IAA homeostasis.

Overall, this work supports differential and developmental stage-specific contributions of the UGT84B1, UGT74D1 and UGT76E3456 glycosyltransferases to IAA homeostasis by mediating IAA and oxIAA glycosylation (Figure 6). Our data indicate that IAA homeostasis is redundantly controlled by UGT84B1, UGT74D1 and UGT76E3456 through IAA and oxIAA glycosylation. The identification of putative additional IAA and oxIAA glycosyltransferases, along with the genetic and biochemical analysis of multiple mutants, will advance our understanding of the contribution of UGTs to IAA homeostasis and plant development.

**Figure 6.**
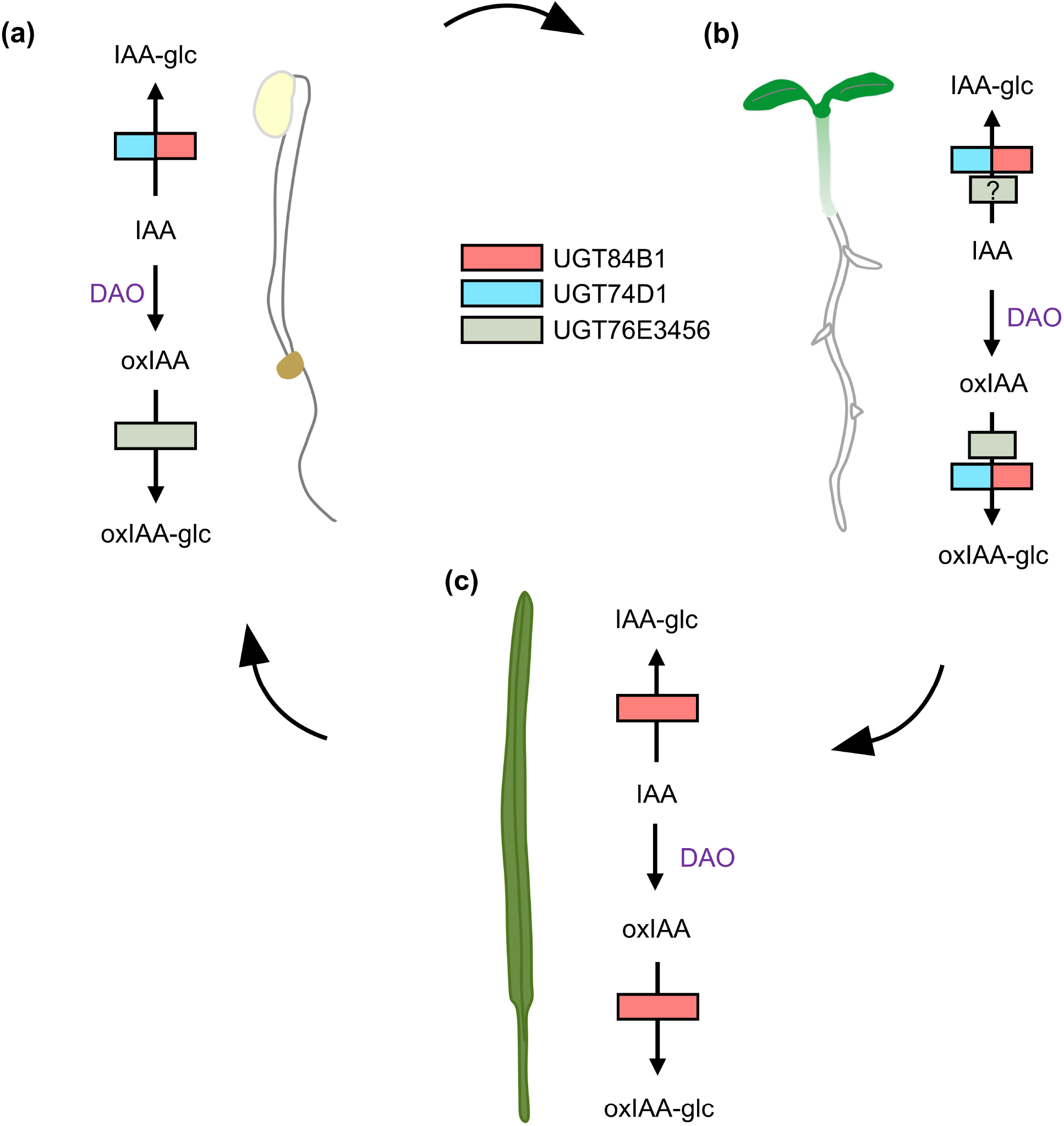
Proposed model of action of UGTs in IAA metabolism at different developmental stages. Simplified IAA inactivation pathways in etiolated seedlings (a), light-grown seedlings (b), and siliques (c). The major players in IAA and oxIAA glycosylation at each stage are indicated by coloured boxes. Smaller boxes indicate likely minor contributions by the UGTs indicated. DAO: DIOXYGENASE FOR AUXIN OXIDATION.

## MATERIALS AND METHODS

### Plant material, culture conditions and IAA treatments

All *Arabidopsis thaliana* plants studied in this work were homozygous for the mutations indicated. The Nottingham Arabidopsis Stock Centre provided seeds for the wild-type accession Col-0 (N1092), as well as seeds of the *ugt76e5* (SALK_006783; N25012), *ugt76e6* (SALK_200519; N687668), and *ugt74d1* (SALK_004870; N504870) mutants. The presence and positions of all T-DNA insertions were confirmed by PCR amplification using gene-specific primers and the LbB1.3 primer (Supplemental Table 1).

Seeds were surface sterilized with bleach solution (40% vol/vol commercial bleach in dH_2_O and 0.002% Triton X-100) for 8 min and then washed four times with sterile deionized water. Seeds were stratified for 3 days and sown under sterile conditions on petri dishes containing half-strength Murashige & Skoog salt mixture (Duchefa, M0221), 1% sucrose, 0.05% MES hydrate (Sigma, M2933) and 0.8% plant agar (Duchefa, P1001), at pH 5.7. Flowering plants were grown in pots containing a 3:1 mixture of organic soil and vermiculite. All plants were grown in long-day conditions (16h light, 8h dark) at 22 ± 1°C under cool white fluorescent light (150 μmol photons m^−2^ s^−1^). IAA treatments and feeding experiments with ^13^C_6_-labelled IAA were performed on 7-day-old seedlings. Seedlings were incubated in half-strength liquid MS media with and without 1 μM of IAA for 1, 2 and 4 hours, or with and without 1 μM of [^13^C_6_]IAA for 12 hours, under gentle shaking and in darkness. The 12-hour timepoint was chosen because we had previously observed a peak of IAA *de novo* glycosylation in Col-0 seedlings after 12 hours of feeding with [^13^C_6_]IAA (Porco *et al.*, 2016).

### Plant phenotyping

For root phenotyping, plants were grown vertically in square petri dishes and plates were imaged using Epson Perfection V600 Photo scanners. For skotomorphogenesis experiments, seeds were stratified at 4°C for 3 days, and then transferred to light at 22 ± 1°C for 8 hours to induce germination. Plates were then transferred to darkness and incubated at 22 ± 1°C for five days. To simulate shade conditions, plants were grown normally for four days and then transferred to darkness for five days. Lengths of the roots and hypocotyls were measured using FIJI software (Schindelin *et al.*, 2012).

### Plasmid construction and plant transformation

The CRISPR/Cas9-based vector for the knockout of the *UGT84B1* gene was constructed using the GreenGate system (Lampropoulos *et al.*, 2013) as described in (Capovilla *et al.*, 2017). Four guide RNAs (gRNAs) targeting the *UGT84B1* coding sequence (Table S1 and Supplemental Figure 14) were designed using CRISPR-P (http://crispr.hzau.edu.cn/CRISPR/). Two supermodules were first created by assembling GreenGate modules into the intermediate plasmid vectors pGGM000 and pGGN000. The M intermediate vector resulted from assembly of the modules A: EC1.2enhancer-EC1.1promoter; B: *Arabidopsis thaliana* codon-optimized Cas9; C: rbcs terminator; D: gRNA sg84b1.4; E: gRNA sg84b1.1; and FH-adapter (pGGG001). The N intermediate vector resulted from assembly of the HA-adapter (pGGG002); A: UBQ10 promoter (pGGA006); B: mCherry; C: rbcs terminator; D: gRNA sg84b1.2; E: gRNA sg84b1.3; F: hygromycin B phosphotransferase (pGGF005). The M and N supermodules were then combined into the destination vector pGGZ003 to create the final construct. The gRNAs were generated using the primers listed in Supplemental Table 1 and cloned into the D and E modules by digestion-ligation (Lampropoulos *et al.*, 2013). The mCherry sequence was amplified from the pGGC015 plasmid using the mCherry-BasI primers (Supplemental Table 1) and cloned into a B module by digestion-ligation (Lampropoulos *et al.*, 2013). The pGGA006, pGGC015, pGGG001, pGGG002, pGGM000 and pGGN000 plasmids were purchased from Addgene. The B:Cas9, C:rbcs terminator, pGGF005 and pGGZ003 were kindly provided by Prof. Markus Schmid. The A module containing the previously reported EC1.2enhancer-EC1.1promoter construct (Wang *et al.*, 2015) was kindly provided by Dr. Wei Wang. The integrity of the insert sequences in all modules was confirmed by Sanger sequencing. The correct assembly of the supermodules in the intermediate and destination vectors was confirmed by restriction analysis.

To construct the *35S_pro_:UGT76E5*, the *UGT76E5* transcription unit, from the ATG to the stop codon, was amplified from Col-0 cDNA using Q5 High-Fidelity DNA Polymerase (NEB), as recommended by the manufacturer. The oligonucleotides attB_UGT76E5_F and attB_UGT76E5_R, which contained *att*B sites at their 5’ ends, were used (Supplemental Table 1). PCR product of the expected size was purified from the agarose gel after electrophoresis using the Monarch DNA Gel Extraction Kit (NEB), and cloned into the pDONR207 vector (Invitrogen) by following the Gateway BP Clonase II Enzyme Mix (Thermo Fisher) protocol. Chemically competent *Escherichia coli* DH5α cells were transformed by the heat-shock method (Dagert & Ehrlich, 1979), and the sequence integrity of the insert carried by transformants was verified by Sanger sequencing. The insert in pDONR207 was subcloned into the pMDC32 Gateway-compatible destination vector (Curtis & Grossniklaus, 2003) via an LR Clonase II (Thermo Fisher) reaction. Seven independent transgenic lines were obtained without any morphological phenotype and high *UGT76E5* expression levels (Supplementary Figure 4). Line #1 was chosen for the metabolic analysis.

To create the CRISPR/Cas9 construct to knock out *UGT76E3*, *UGT76E4* and *UGT76E5*, the pKI1.1R plasmid was used following the protocol described in Tsutsui and Higashiyama (2017). Briefly, the circular pKI1.1R plasmid was linearized by incubating 1.5 μg of the purified plasmid with the AarI restriction enzyme for 10 hours, and then dephosphorylated using FastAP (Thermo Fisher). A target-specific gRNA was designed using CRISPR-P 2.0 (http://crispr.hzau.edu.cn/CRISPR2/). Oligonucleotides harbouring the gRNA target (sgRNA_UGT76E345_F and sgRNA_UGT76E345_R; Supplementary Table 1) were hybridized by slow cooling from 95-25°C and then phosphorylated using T4 Polynucleotide Kinase (NEB). The digested plasmid and the hybridized oligonucleotides were ligated using T4 Ligase (NEB) and then transformed into *Escherichia coli* DH5α competent cells. The sequence integrity of inserts carried by transformants was verified by Sanger sequencing.

All constructs were mobilized into *Agrobacterium tumefaciens* GV3101 (C58C1 Rif^R^) cells. The CRISPR-UGT84B1 and the *35S_pro_:UGT76E5* constructs were used to transform Col-0 and CRISPR-UGT76E345 was used to transform *ugt76e6* plants by the floral dip method (Clogh and Bent, 1998). T_1_ transgenic plants were selected on plates supplemented with 15 mg/L hygromycin B (Invitrogen).

### RNA isolation, cDNA synthesis, and qRT-PCR

For qRT-PCR, total RNA was isolated using an RNeasy Plant Mini Kit (Qiagen). DNA was removed using a TURBO DNA-free Kit (Invitrogen). First-strand cDNA synthesis was performed using an iScript cDNA Synthesis Kit (BioRad) following the manufacturer’s instructions. *ACTIN2* was used as an internal control in relative expression analyses. Three biological replicates were analysed in triplicate. qPCR reactions were performed in a 10-μl volume containing 5 μl of LightCycler 480 SYBR Green I Master (Roche), 4 μl of the corresponding primer pair (1.5 μM each), and 1 μl of cDNA template. Quantification of relative gene expression was performed using the comparative C_T_ method (2-^ΔΔCt^) (Schmittgen & Livak, 2008) on a CFX96 Real-Time System (BioRad). Primers used are listed in Supplemental Table 1.

### Bioinformatics analyses

To identify UGT76E5 paralogs, BLASTP searches (Altschul *et al.*, 1997) were performed at NCBI using the refseq_protein database and default parameters. Proteins with a percentage of sequence identity higher than 70% were selected, aligned using ClustalW (Thompson *et al.*, 1994), and shaded with BOXSHADE3.21 (https://embnet.vital-it.ch/software/BOX_form.html). A phylogenetic tree was constructed by the neighbour-joining clustering method, inferred from 1000 replicates, with MEGA X (Kumar *et al.*, 2018) using default parameters (model: Poisson; rates among sites: uniform rates; gaps/missing data treatment: pairwise deletion). Auxin-related genes co-expressed with *UGT76E3*, *UGT76E4*, *UGT76E5*, and *UGT76E6* were retrieved from the ATTED-II database (Obayashi *et al.*, 2018).

### IAA metabolite profiling

Extraction and purification of the targeted compounds (IAA, oxIAA, IAA-Asp, IAA-Glu IAA-Glc, oxIAA-Glc, both unlabelled and [^13^C_6_] labelled compounds) were performed according to (Novák *et al.*, 2012), with slight modifications. Briefly, 10 mg of frozen material per sample was homogenized using a bead mill (27 Hz, 10 min, 4°C; MixerMill, Retsch GmbH, Haan, Germany) and extracted in 1 ml of 50 mM sodium phosphate buffer containing 1% sodium diethyldithiocarbamate and a mixture of deuterium/nitrogen isotopically labelled internal standards ([^2^H_5_]IAA, ([^2^H_4_]oxIAA, [^15^N,^2^H_5_]IAA-Asp, [^15^N,^2^H_5_]IAA-Glu, Olchemim, Olomouc, Czech Republic),. After centrifugation (20 000 *g*, 15 min, 4°C), the supernatant was transferred into new Eppendorf tubes. The pH was then adjusted to 2.5 with 1 M HCl and samples were immediately applied to preconditioned solid‐phase extraction columns (Oasis HLB, 30 mg of 1 ml; Waters Inc., Milford, MA, USA). After sample application, each column was rinsed with 2 ml 5% methanol. Compounds of interest were then eluted with 2 ml 80% methanol. UHPLC-MS/MS analysis was performed according to the method described in (Pěnčík *et al.*, 2018), using an LC‐MS/MS system consisting of a 1290 Infinity Binary LC System coupled to a 6490 Triple Quad LC/MS System with Jet Stream and Dual Ion Funnel technologies (Agilent Technologies, Santa Clara, CA, USA).

### Accession Numbers

*ACTIN2* (At3g18780), *UGT74D1* (At2g31750), *UGT76E3* (At3g46700), *UGT76E4* (At3g46690), *UGT76E5* (At3g46720), *UGT76E6* (At3g46680), *UGT84B1* (At2g23260).

## SUPPLEMENTARY MATERIAL

**Supplementary Figure 1. Morphological phenotypes** of *ugt84b1*, *ugt74d1 ugt76e5*, and *ugt76e3456* plants.

**Supplementary Figure 2.** Tissue-specific profiling of IAA catabolites in the *ugt84b1* mutant.

**Supplementary Figure 3.** *UGT76E3, UGT76E4, UGT76E5, and UGT76E6* are co-expressed with auxin-related genes.

**Supplementary Figure 4.** qRT-PCR analysis of the relative expression of *UGT76E5* in seven independent transformants carrying the *35S_pro_:UGT76E5* transgene.

**Supplementary Figure 5**. Phylogenetic analysis of the UGT76 family.

**Supplementary Figure 6.** Alignment of the amino acid sequences of the Arabidopsis proteins UGT76E3, UGT76E4, UGT76E5 and UGT76E6.

**Supplementary Figure 7.** The *UGT76E3*, *UGT76E4*, *UGT76E5*, and *UGT76E6* genes are clustered together **on** Arabidopsis chromosome 3.

**Supplementary Figure 8.** CRISPR-based approach to knock out the entire UGT76E3456 subfamily.

**Supplementary Figure 9.** Detailed information about the *UGT76E3* gene editing.

**Supplementary Figure 10.** Detailed information about the *UGT76E4* gene editing.

**Supplementary Figure 11.** Detailed information about the *UGT76E5* gene editing.

**Supplementary Figure 12.** *UGT74D1* expression pattern and molecular characterization of the *ugt74d1* mutant.

**Supplementary Figure 13.** Steady-state levels of IAA metabolites in *ugt84b1*, *ugt74d1, ugt76e3456* and *35S_pro_:UGT76E5* plants.

**Supplementary Figure 14.** Sequence information for the *UGT84B1* gene editing.

**Supplementary Table 1**. Primer sets used in this work.

## ACKNOWLEDGMENTS

Research in the laboratory of Karin Ljung is supported by grants from the Swedish Foundation for Strategic Research (Vinnova), the Knut and Alice Wallenberg Foundation (KAW), the Swedish research councils VR and Formas, and Carl Tryggers Stiftelse för Vetenskaplig Forskning. E.M.-B. (JCK-1811) and R.C.-S. (JCK-1111) held postdoctoral fellowships from Kempestiftelserna. We also acknowledge the Swedish Metabolomics Centre (http://www.swedishmetabolomicscentre.se/) for access to instrumentation.

## AUTHOR CONTRIBUTIONS

E.M.-B., R.C.-S. and K.L. conceived and designed the research; E.M.-B. and R.C.-S. performed most of the experiments; J.Š. performed the hormone analyses. E.M.-B. wrote the manuscript draft and prepared the figures and tables; E.M.-B. and R.C.-S. wrote the manuscript with input from all authors. This research was supported by funds to K.L.

