## Supplementary Table 1 for "Redefining the roles of UDP-glycosyltransferases in auxin metabolism and homeostasis during plant development"

**Supplemental Table 1**. Primer sets used in this work

| Purpose | Oligonucleotide name(s) | Oligonucleotide sequences (5' → 3') | |
|---|---|---|---|
| | | Forward primer (F) | Reverse primer (R) |
| Genotyping | SALK_006783 | TTCACAATGCATCCTCTTTCC | CAAACAAGTGACGTGTCAACC |
| | SALK_200519 | TCTATCTGTGGTGCCCATTTC | TTGTCACACAAGGAACACGAG |
| | SALK_004870 | ACATACTGGCTAGGCATGTGG | TAGTCGGCTGGTCGCTATAAG |
| | LBb1.3 | | ATTTTGCCGATTTCGGAAC |
| | UGT76E3_tar_ | TGACTTTGCAGAGCATGACG | AAACAAGGGTTCTCTCCCTCA |
| | UGT76E4_tar_ | AATGACCCATTTCGAAAGCA | ATGGCTGCACCTTCTTCATC |
| | UGT76E5_tar_ | TACCGATTCTGCAACCCTTC | ACAAGGGCTCTCTCCCTCAT |
| | UGT84B1_tar_ | TCGTGTTCTTCTCCGATGGT | ATTCCGTGTCGAAGATCCAC |
| qRT-PCR | q_ACT2 | GCACCCTGTTCTTCTTACCG | AACCCTCGTAGATTGGCACA |
| | q_UGT76E5 | AAGCAAACTCAACGCCGAGA | ATGTTTTGCACTTCAGGATCTTTCA |
| | q_UGT74D1 | TCTCTAAAAACGTCAACGTCACA | CGGAGGATGGAGTTGTGG |
| | q_UGT84B1 | TGTAGGTTCTTTGATGATAACGA | CACATGTCCTAATGGTAACAC |
| Cloning | attB_UGT76E5 | GGGGACAAGTTTGTACAAAAAAGCAGGCTATGGAGAAAA ATGCAGAGAAGAAA | GGGGACCACTTTGTACAAGAAAGCTGGGTTCAAGTATTT CTATACTCTGCCT |
| | sgRNA_UGT84B1.1 | GAGAAGTGAGCACCACCAGAGTTTTAGAGCTATGCTG | TCTGGTGGTGCTCACTTCTCAATCACTACTTCGACTC |
| | sgRNA_UGT84B1.2 | GGTAATTCCAATTGGTCCTCGTTTTAGAGCTATGCTG | GAGGACCAATTGGAATTACCAATCACTACTTCGACTC |
| | sgRNA_UGT84B1.3 | GTTCTTCAGCGCCTTCGCTAGTTTTAGAGCTATGCTG | TAGCGAAGGCGCTGAAGAACAATCACTACTTCGACTC |
| | sgRNA_UGT84B1.4 | GGTAGCGTACCCTAGCTGGAGTTTTAGAGCTATGCTG | TCCAGCTAGGGTACGCTACCAATCACTACTTCGACTC |
| | mCherry-BasI | AACAGGTCTCAAACAATGGTGAGCAAGGGCGAGG | AACAGGTCTCAAGCCTTACTTGTACAGCTCGTCCATGC |
| | sgRNA_UGT76E345 | AATGGTAATGTGAAGAGGGCCTAA | AAACTTAGGCCCTCTTCACATTAC |
