## Supplementary Figures for "Redefining the roles of UDP-glycosyltransferases in auxin metabolism and homeostasis during plant development"

### Slide 1
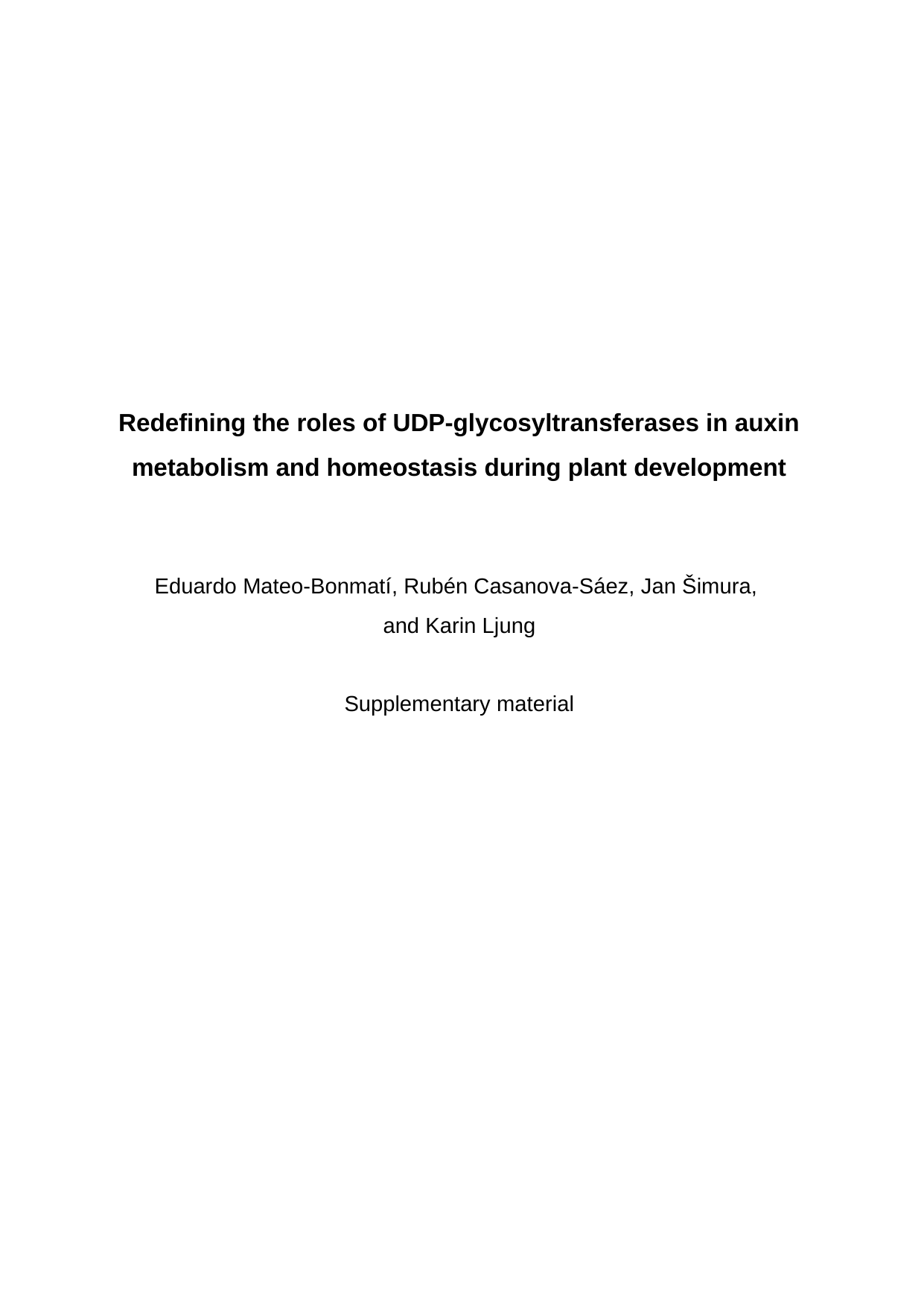

#### Redefining the roles of UDP-glycosyltransferases in auxin metabolism and homeostasis during plant development Eduardo Mateo-Bonmatí, Rubén Casanova-Sáez, Jan Šimura, and Karin Ljung Supplementary material

### Slide 2
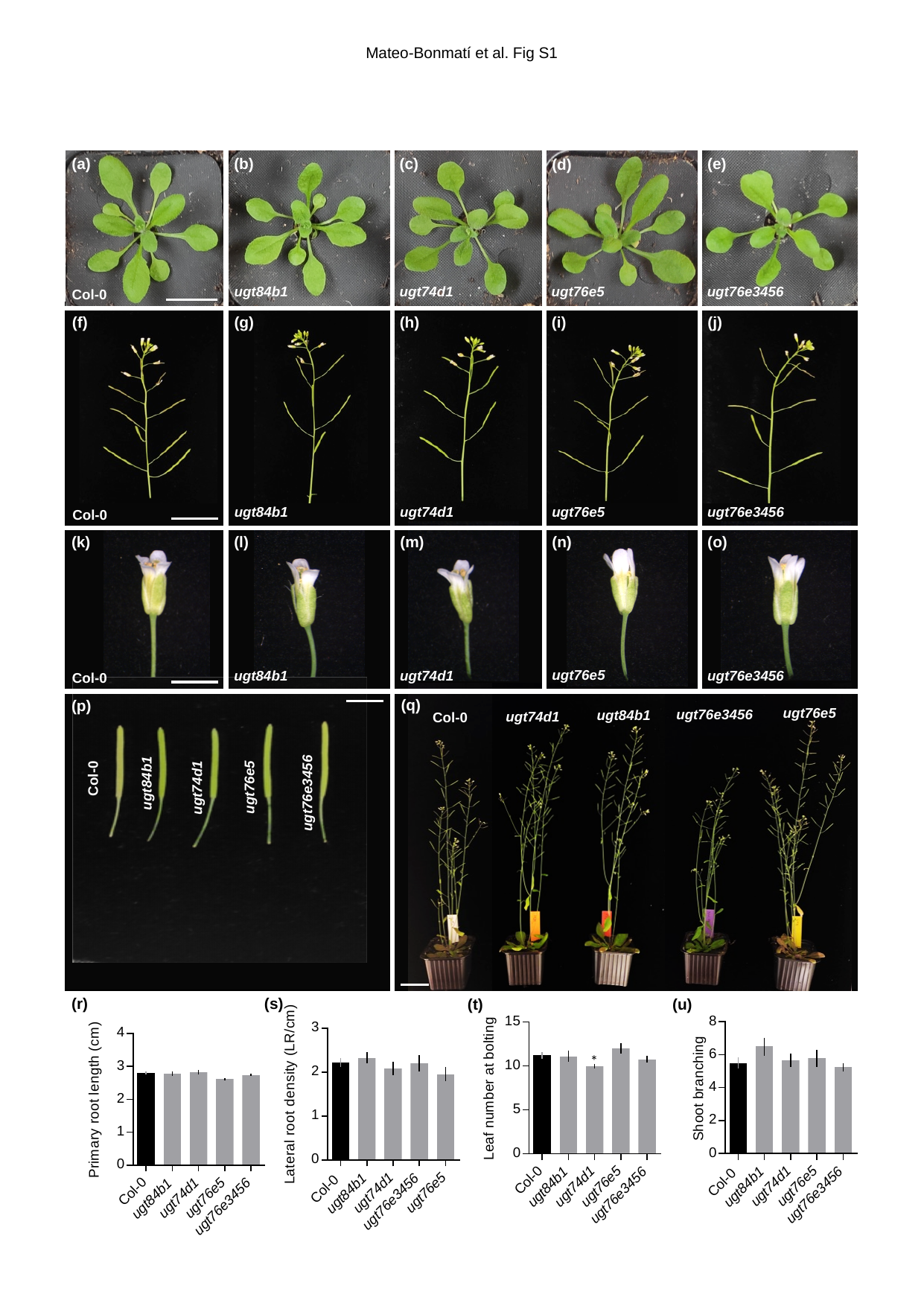

Mateo-Bonmatí et al. Fig S1
(a)
(b)
(c)
(e)
(d)
ugt76e5
ugt84b1
ugt76e3456
ugt74d1
Col-0
(g)
(f)
(j)
(i)
(h)
ugt76e5
ugt84b1
ugt74d1
ugt76e3456
Col-0
(o)
(l)
(k)
(n)
(m)
ugt76e5
ugt74d1
ugt76e3456
ugt84b1
Col-0
Col-0
ugt84b1
ugt74d1
ugt76e5
ugt76e3456
ugt76e5
ugt76e3456
ugt84b1
ugt74d1
Col-0
(q)
(p)
(r)
(s)
(t)
(u)
*
*

### Slide 3
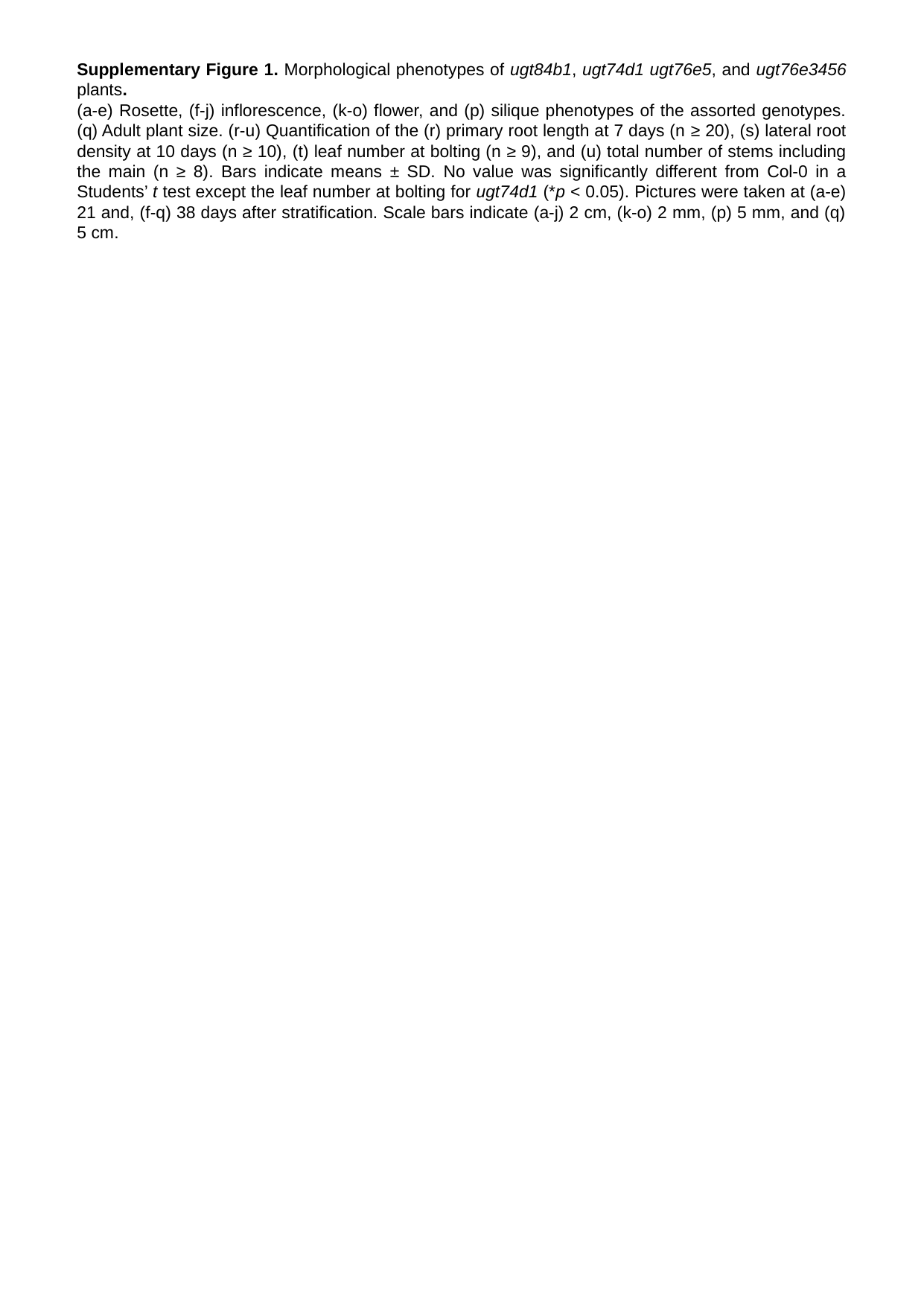

Supplementary Figure 1. Morphological phenotypes of ugt84b1, ugt74d1 ugt76e5, and ugt76e3456 plants.
(a-e) Rosette, (f-j) inflorescence, (k-o) flower, and (p) silique phenotypes of the assorted genotypes. (q) Adult plant size. (r-u) Quantification of the (r) primary root length at 7 days (n ≥ 20), (s) lateral root density at 10 days (n ≥ 10), (t) leaf number at bolting (n ≥ 9), and (u) total number of stems including the main (n ≥ 8). Bars indicate means ± SD. No value was significantly different from Col-0 in a Students’ t test except the leaf number at bolting for ugt74d1 (*p < 0.05). Pictures were taken at (a-e) 21 and, (f-q) 38 days after stratification. Scale bars indicate (a-j) 2 cm, (k-o) 2 mm, (p) 5 mm, and (q) 5 cm.

### Slide 4
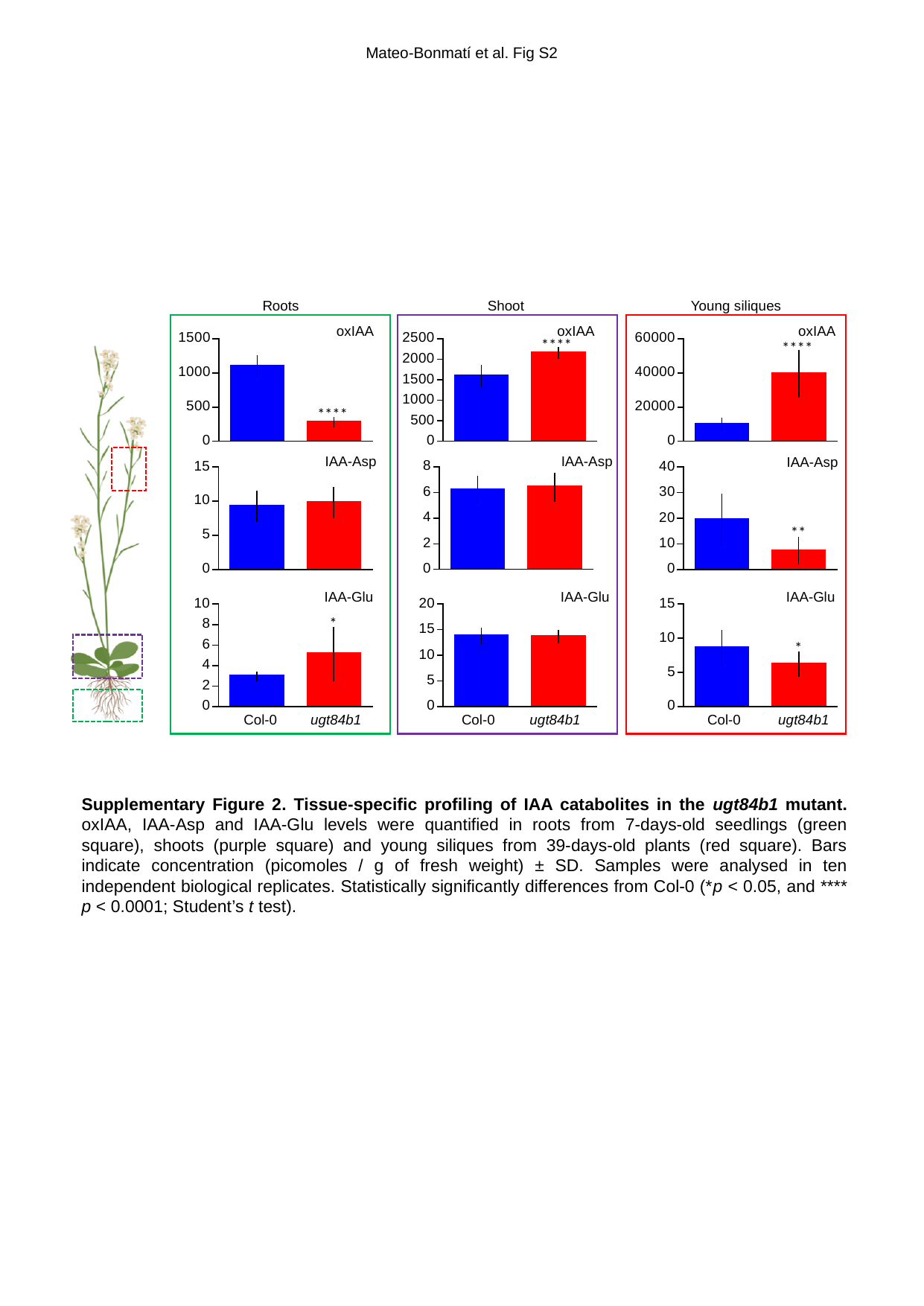

Mateo-Bonmatí et al. Fig S2
Roots
Shoot
Young siliques
oxIAA
oxIAA
oxIAA
****
****
****
IAA-Asp
IAA-Asp
IAA-Asp
**
IAA-Glu
IAA-Glu
IAA-Glu
*
*
Col-0
ugt84b1
Col-0
ugt84b1
Col-0
ugt84b1
Supplementary Figure 2. Tissue-specific profiling of IAA catabolites in the ugt84b1 mutant. oxIAA, IAA-Asp and IAA-Glu levels were quantified in roots from 7-days-old seedlings (green square), shoots (purple square) and young siliques from 39-days-old plants (red square). Bars indicate concentration (picomoles / g of fresh weight) ± SD. Samples were analysed in ten independent biological replicates. Statistically significantly differences from Col-0 (*p < 0.05, and **** p < 0.0001; Student’s t test).

### Slide 5
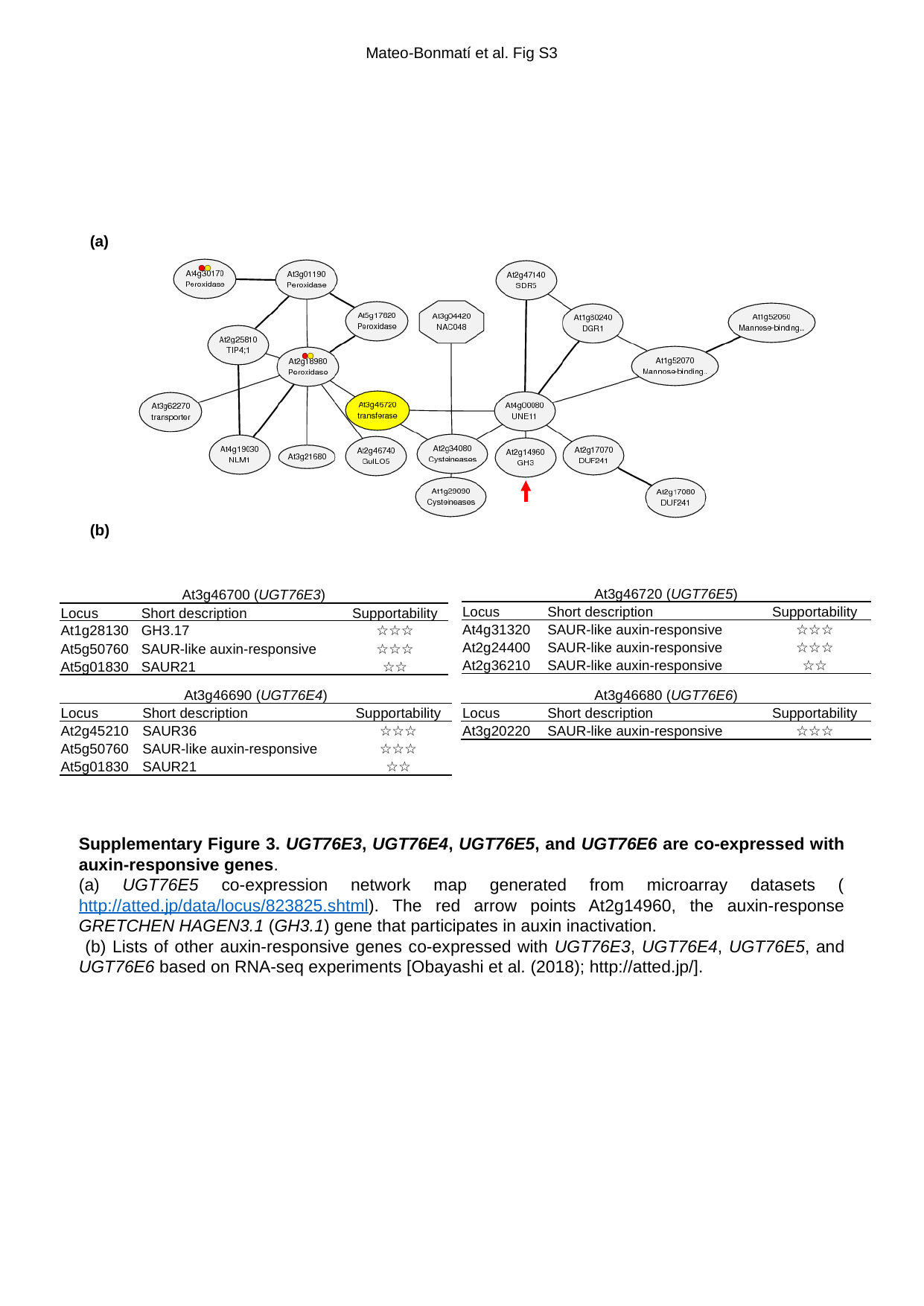

Mateo-Bonmatí et al. Fig S3
(a)
(b)
| At3g46720 (UGT76E5) | | |
| --- | --- | --- |
| Locus | Short description | Supportability |
| At4g31320 | SAUR-like auxin-responsive | ☆☆☆ |
| At2g24400 | SAUR-like auxin-responsive | ☆☆☆ |
| At2g36210 | SAUR-like auxin-responsive | ☆☆ |
| At3g46700 (UGT76E3) | | |
| --- | --- | --- |
| Locus | Short description | Supportability |
| At1g28130 | GH3.17 | ☆☆☆ |
| At5g50760 | SAUR-like auxin-responsive | ☆☆☆ |
| At5g01830 | SAUR21 | ☆☆ |
| At3g46690 (UGT76E4) | | |
| --- | --- | --- |
| Locus | Short description | Supportability |
| At2g45210 | SAUR36 | ☆☆☆ |
| At5g50760 | SAUR-like auxin-responsive | ☆☆☆ |
| At5g01830 | SAUR21 | ☆☆ |
| At3g46680 (UGT76E6) | | |
| --- | --- | --- |
| Locus | Short description | Supportability |
| At3g20220 | SAUR-like auxin-responsive | ☆☆☆ |
Supplementary Figure 3. UGT76E3, UGT76E4, UGT76E5, and UGT76E6 are co-expressed with auxin-responsive genes.
(a) UGT76E5 co-expression network map generated from microarray datasets (http://atted.jp/data/locus/823825.shtml). The red arrow points At2g14960, the auxin-response GRETCHEN HAGEN3.1 (GH3.1) gene that participates in auxin inactivation.
 (b) Lists of other auxin-responsive genes co-expressed with UGT76E3, UGT76E4, UGT76E5, and UGT76E6 based on RNA-seq experiments [Obayashi et al. (2018); http://atted.jp/].

### Slide 6
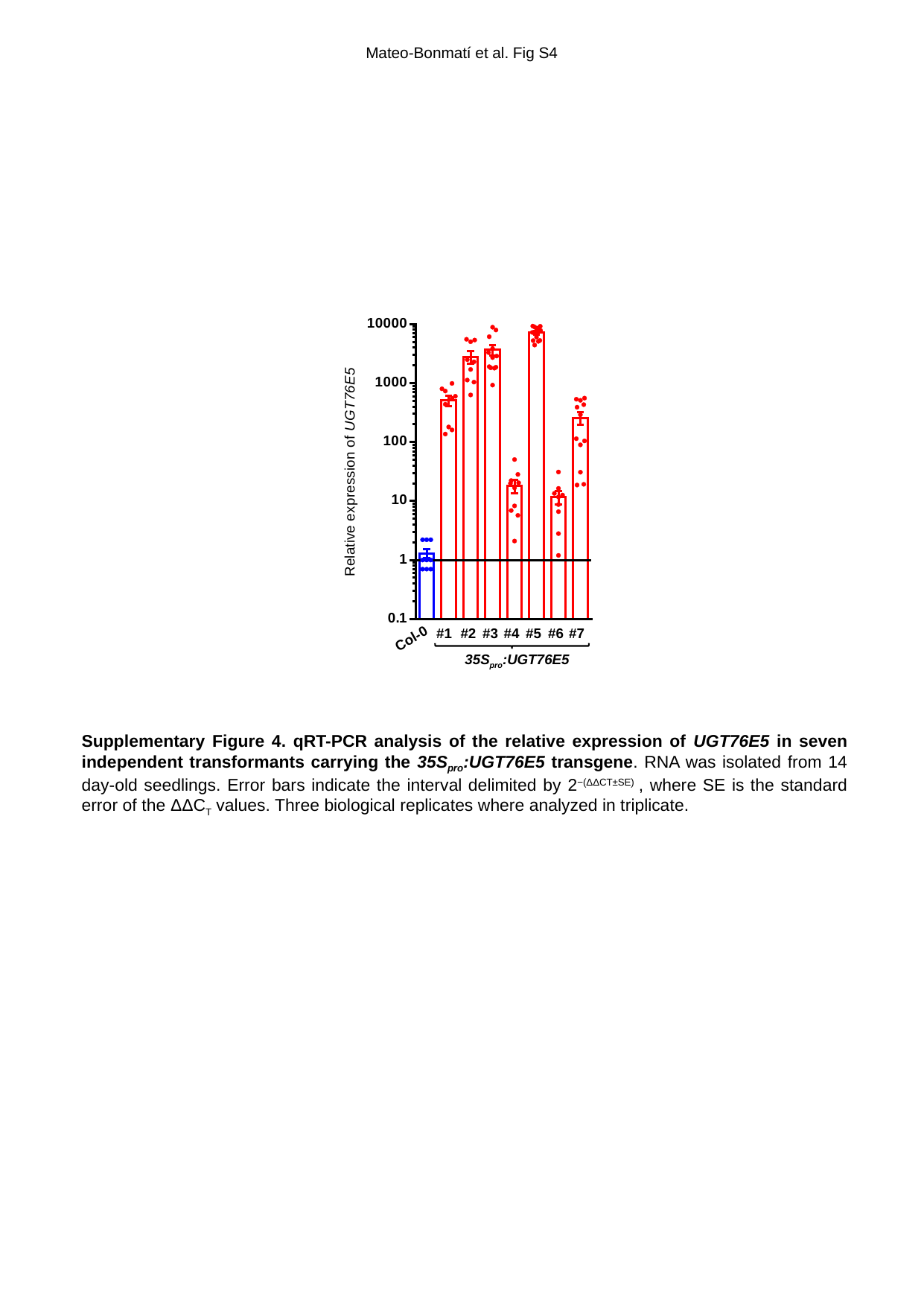

Mateo-Bonmatí et al. Fig S4
#1
#2
#3
#4
#5
#6
#7
Col-0
35S_pro_:UGT76E5
Supplementary Figure 4. qRT-PCR analysis of the relative expression of UGT76E5 in seven independent transformants carrying the 35S_pro_:UGT76E5 transgene. RNA was isolated from 14 day-old seedlings. Error bars indicate the interval delimited by 2^−(ΔΔCT±SE)^ , where SE is the standard error of the ΔΔC_T_ values. Three biological replicates where analyzed in triplicate.

### Slide 7
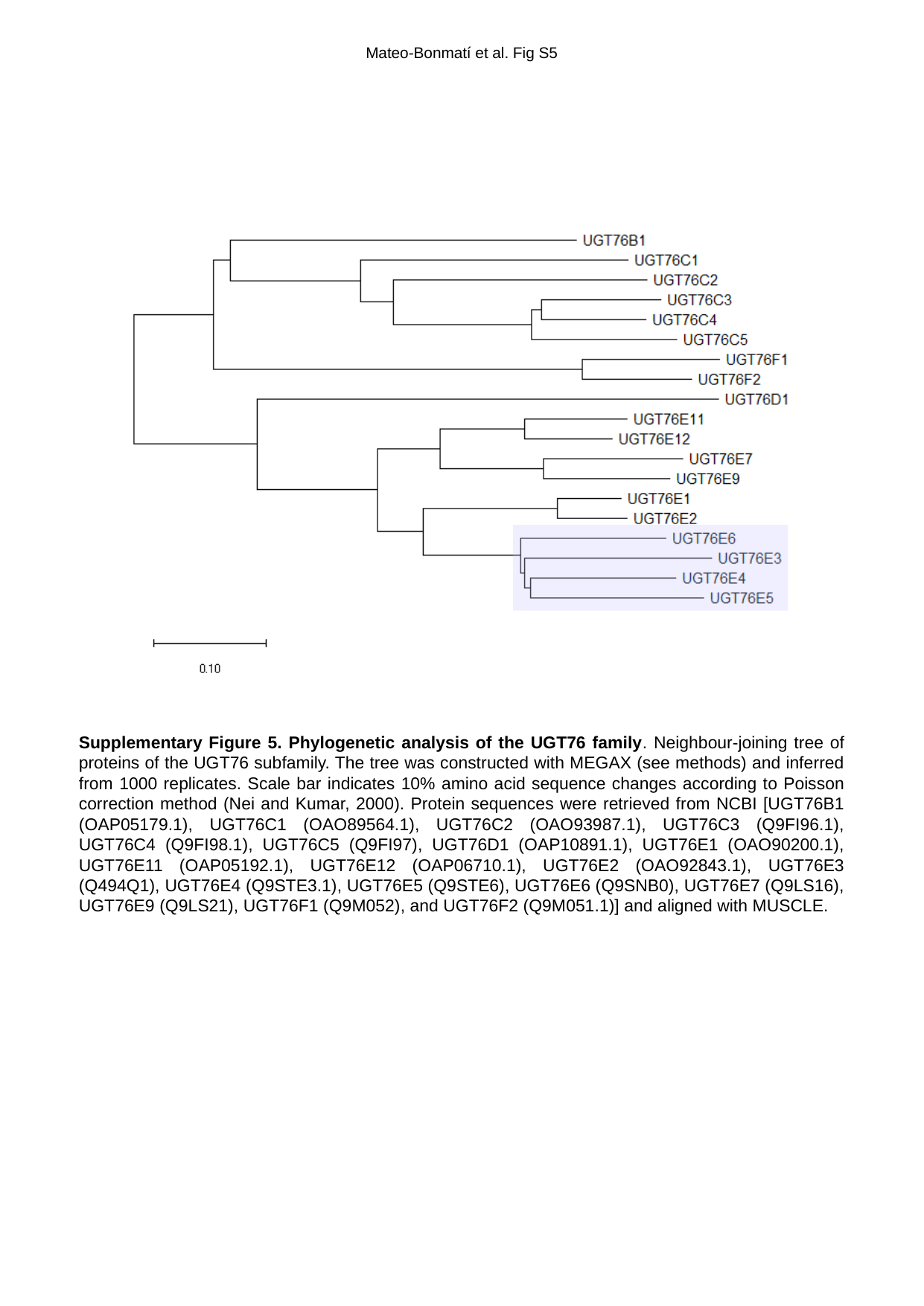

Mateo-Bonmatí et al. Fig S5
Supplementary Figure 5. Phylogenetic analysis of the UGT76 family. Neighbour-joining tree of proteins of the UGT76 subfamily. The tree was constructed with MEGAX (see methods) and inferred from 1000 replicates. Scale bar indicates 10% amino acid sequence changes according to Poisson correction method (Nei and Kumar, 2000). Protein sequences were retrieved from NCBI [UGT76B1 (OAP05179.1), UGT76C1 (OAO89564.1), UGT76C2 (OAO93987.1), UGT76C3 (Q9FI96.1), UGT76C4 (Q9FI98.1), UGT76C5 (Q9FI97), UGT76D1 (OAP10891.1), UGT76E1 (OAO90200.1), UGT76E11 (OAP05192.1), UGT76E12 (OAP06710.1), UGT76E2 (OAO92843.1), UGT76E3 (Q494Q1), UGT76E4 (Q9STE3.1), UGT76E5 (Q9STE6), UGT76E6 (Q9SNB0), UGT76E7 (Q9LS16), UGT76E9 (Q9LS21), UGT76F1 (Q9M052), and UGT76F2 (Q9M051.1)] and aligned with MUSCLE.

### Slide 8
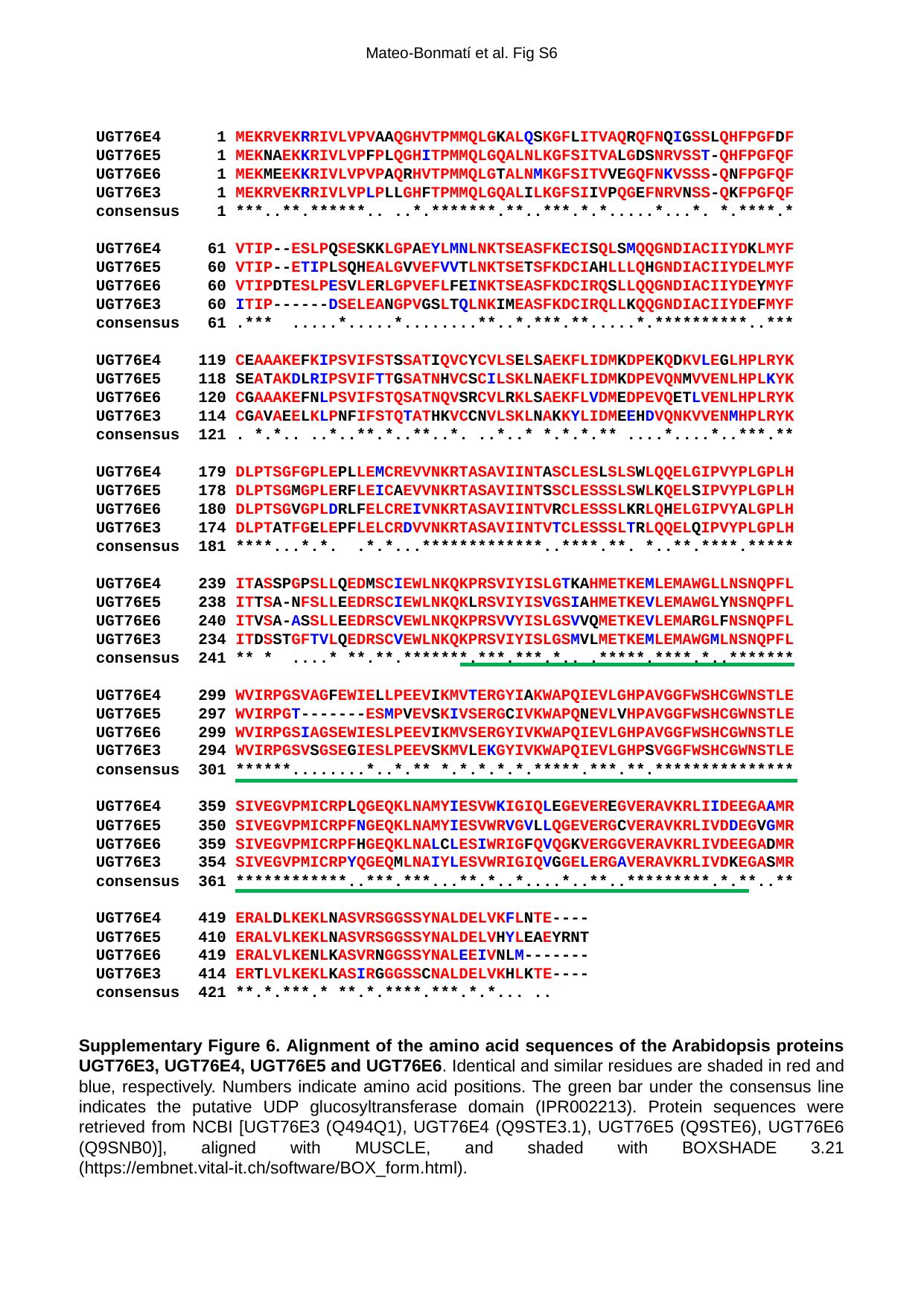

Mateo-Bonmatí et al. Fig S6
UGT76E4 1 MEKRVEKRRIVLVPVAAQGHVTPMMQLGKALQSKGFLITVAQRQFNQIGSSLQHFPGFDF
UGT76E5 1 MEKNAEKKRIVLVPFPLQGHITPMMQLGQALNLKGFSITVALGDSNRVSST-QHFPGFQF
UGT76E6 1 MEKMEEKKRIVLVPVPAQRHVTPMMQLGTALNMKGFSITVVEGQFNKVSSS-QNFPGFQF
UGT76E3 1 MEKRVEKRRIVLVPLPLLGHFTPMMQLGQALILKGFSIIVPQGEFNRVNSS-QKFPGFQF
consensus 1 ***..**.******.. ..*.*******.**..***.*.*.....*...*. *.****.*
UGT76E4 61 VTIP--ESLPQSESKKLGPAEYLMNLNKTSEASFKECISQLSMQQGNDIACIIYDKLMYF
UGT76E5 60 VTIP--ETIPLSQHEALGVVEFVVTLNKTSETSFKDCIAHLLLQHGNDIACIIYDELMYF
UGT76E6 60 VTIPDTESLPESVLERLGPVEFLFEINKTSEASFKDCIRQSLLQQGNDIACIIYDEYMYF
UGT76E3 60 ITIP------DSELEANGPVGSLTQLNKIMEASFKDCIRQLLKQQGNDIACIIYDEFMYF
consensus 61 .*** .....*.....*........**..*.***.**.....*.**********..***
UGT76E4 119 CEAAAKEFKIPSVIFSTSSATIQVCYCVLSELSAEKFLIDMKDPEKQDKVLEGLHPLRYK
UGT76E5 118 SEATAKDLRIPSVIFTTGSATNHVCSCILSKLNAEKFLIDMKDPEVQNMVVENLHPLKYK
UGT76E6 120 CGAAAKEFNLPSVIFSTQSATNQVSRCVLRKLSAEKFLVDMEDPEVQETLVENLHPLRYK
UGT76E3 114 CGAVAEELKLPNFIFSTQTATHKVCCNVLSKLNAKKYLIDMEEHDVQNKVVENMHPLRYK
consensus 121 . *.*.. ..*..**.*..**..*. ..*..* *.*.*.** ....*....*..***.**
UGT76E4 179 DLPTSGFGPLEPLLEMCREVVNKRTASAVIINTASCLESLSLSWLQQELGIPVYPLGPLH
UGT76E5 178 DLPTSGMGPLERFLEICAEVVNKRTASAVIINTSSCLESSSLSWLKQELSIPVYPLGPLH
UGT76E6 180 DLPTSGVGPLDRLFELCREIVNKRTASAVIINTVRCLESSSLKRLQHELGIPVYALGPLH
UGT76E3 174 DLPTATFGELEPFLELCRDVVNKRTASAVIINTVTCLESSSLTRLQQELQIPVYPLGPLH
consensus 181 ****...*.*. .*.*...*************..****.**. *..**.****.*****
UGT76E4 239 ITASSPGPSLLQEDMSCIEWLNKQKPRSVIYISLGTKAHMETKEMLEMAWGLLNSNQPFL
UGT76E5 238 ITTSA-NFSLLEEDRSCIEWLNKQKLRSVIYISVGSIAHMETKEVLEMAWGLYNSNQPFL
UGT76E6 240 ITVSA-ASSLLEEDRSCVEWLNKQKPRSVVYISLGSVVQMETKEVLEMARGLFNSNQPFL
UGT76E3 234 ITDSSTGFTVLQEDRSCVEWLNKQKPRSVIYISLGSMVLMETKEMLEMAWGMLNSNQPFL
consensus 241 ** * ....* **.**.*******.***.***.*.. .*****.****.*..*******
UGT76E4 299 WVIRPGSVAGFEWIELLPEEVIKMVTERGYIAKWAPQIEVLGHPAVGGFWSHCGWNSTLE
UGT76E5 297 WVIRPGT-------ESMPVEVSKIVSERGCIVKWAPQNEVLVHPAVGGFWSHCGWNSTLE
UGT76E6 299 WVIRPGSIAGSEWIESLPEEVIKMVSERGYIVKWAPQIEVLGHPAVGGFWSHCGWNSTLE
UGT76E3 294 WVIRPGSVSGSEGIESLPEEVSKMVLEKGYIVKWAPQIEVLGHPSVGGFWSHCGWNSTLE
consensus 301 ******........*..*.** *.*.*.*.*.*****.***.**.***************
UGT76E4 359 SIVEGVPMICRPLQGEQKLNAMYIESVWKIGIQLEGEVEREGVERAVKRLIIDEEGAAMR
UGT76E5 350 SIVEGVPMICRPFNGEQKLNAMYIESVWRVGVLLQGEVERGCVERAVKRLIVDDEGVGMR
UGT76E6 359 SIVEGVPMICRPFHGEQKLNALCLESIWRIGFQVQGKVERGGVERAVKRLIVDEEGADMR
UGT76E3 354 SIVEGVPMICRPYQGEQMLNAIYLESVWRIGIQVGGELERGAVERAVKRLIVDKEGASMR
consensus 361 ************..***.***...**.*..*....*..**..*********.*.**..**
UGT76E4 419 ERALDLKEKLNASVRSGGSSYNALDELVKFLNTE----
UGT76E5 410 ERALVLKEKLNASVRSGGSSYNALDELVHYLEAEYRNT
UGT76E6 419 ERALVLKENLKASVRNGGSSYNALEEIVNLM-------
UGT76E3 414 ERTLVLKEKLKASIRGGGSSCNALDELVKHLKTE----
consensus 421 **.*.***.* **.*.****.***.*.*... ..
Supplementary Figure 6. Alignment of the amino acid sequences of the Arabidopsis proteins UGT76E3, UGT76E4, UGT76E5 and UGT76E6. Identical and similar residues are shaded in red and blue, respectively. Numbers indicate amino acid positions. The green bar under the consensus line indicates the putative UDP glucosyltransferase domain (IPR002213). Protein sequences were retrieved from NCBI [UGT76E3 (Q494Q1), UGT76E4 (Q9STE3.1), UGT76E5 (Q9STE6), UGT76E6 (Q9SNB0)], aligned with MUSCLE, and shaded with BOXSHADE 3.21 (https://embnet.vital-it.ch/software/BOX_form.html).

### Slide 9
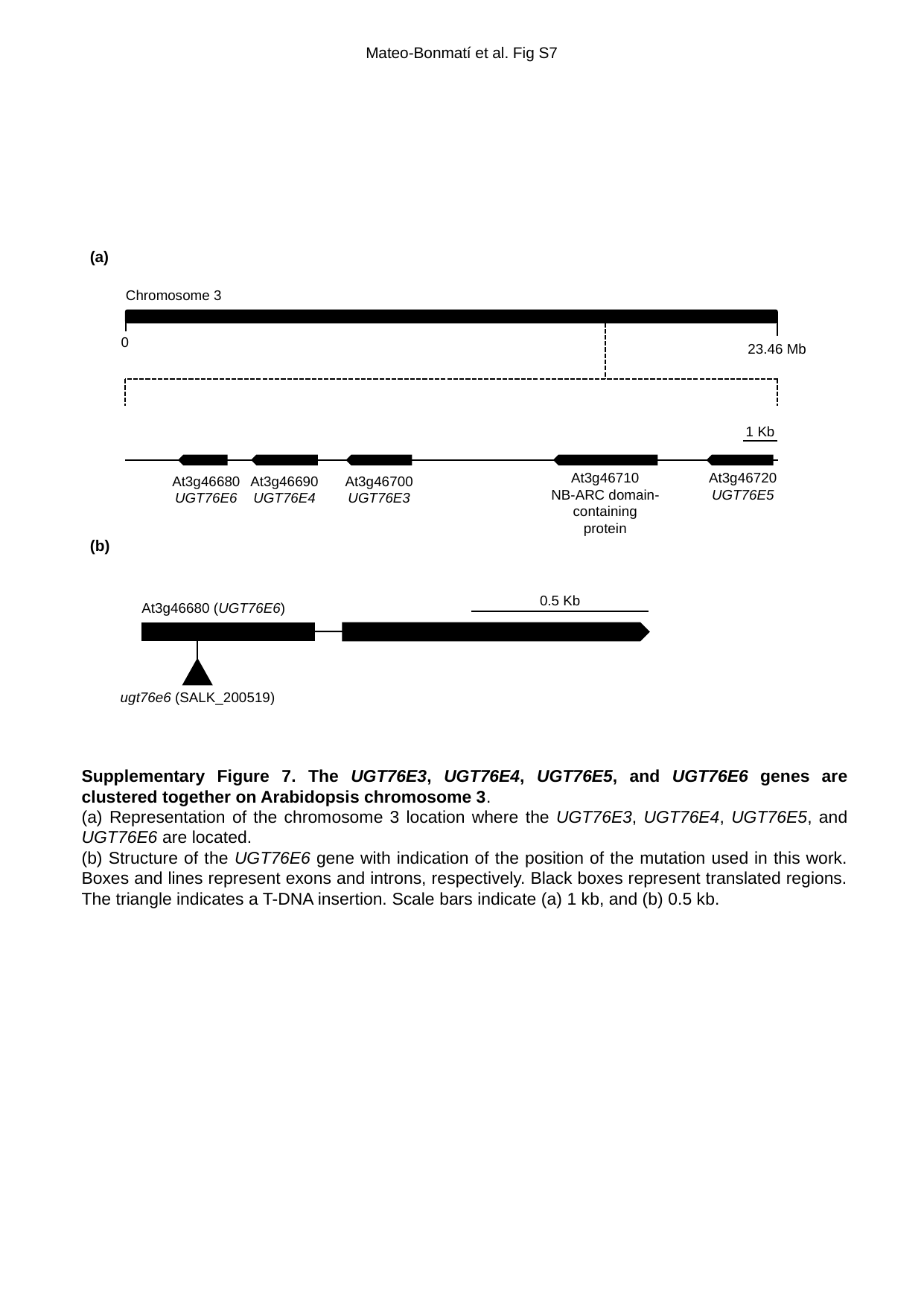

Mateo-Bonmatí et al. Fig S7
(a)
Chromosome 3
0
23.46 Mb
1 Kb
At3g46710
NB-ARC domain-containing protein
At3g46720
UGT76E5
At3g46680
UGT76E6
At3g46700
UGT76E3
At3g46690
UGT76E4
(b)
0.5 Kb
At3g46680 (UGT76E6)
ugt76e6 (SALK_200519)
Supplementary Figure 7. The UGT76E3, UGT76E4, UGT76E5, and UGT76E6 genes are clustered together on Arabidopsis chromosome 3.
(a) Representation of the chromosome 3 location where the UGT76E3, UGT76E4, UGT76E5, and UGT76E6 are located.
(b) Structure of the UGT76E6 gene with indication of the position of the mutation used in this work. Boxes and lines represent exons and introns, respectively. Black boxes represent translated regions. The triangle indicates a T-DNA insertion. Scale bars indicate (a) 1 kb, and (b) 0.5 kb.

### Slide 10
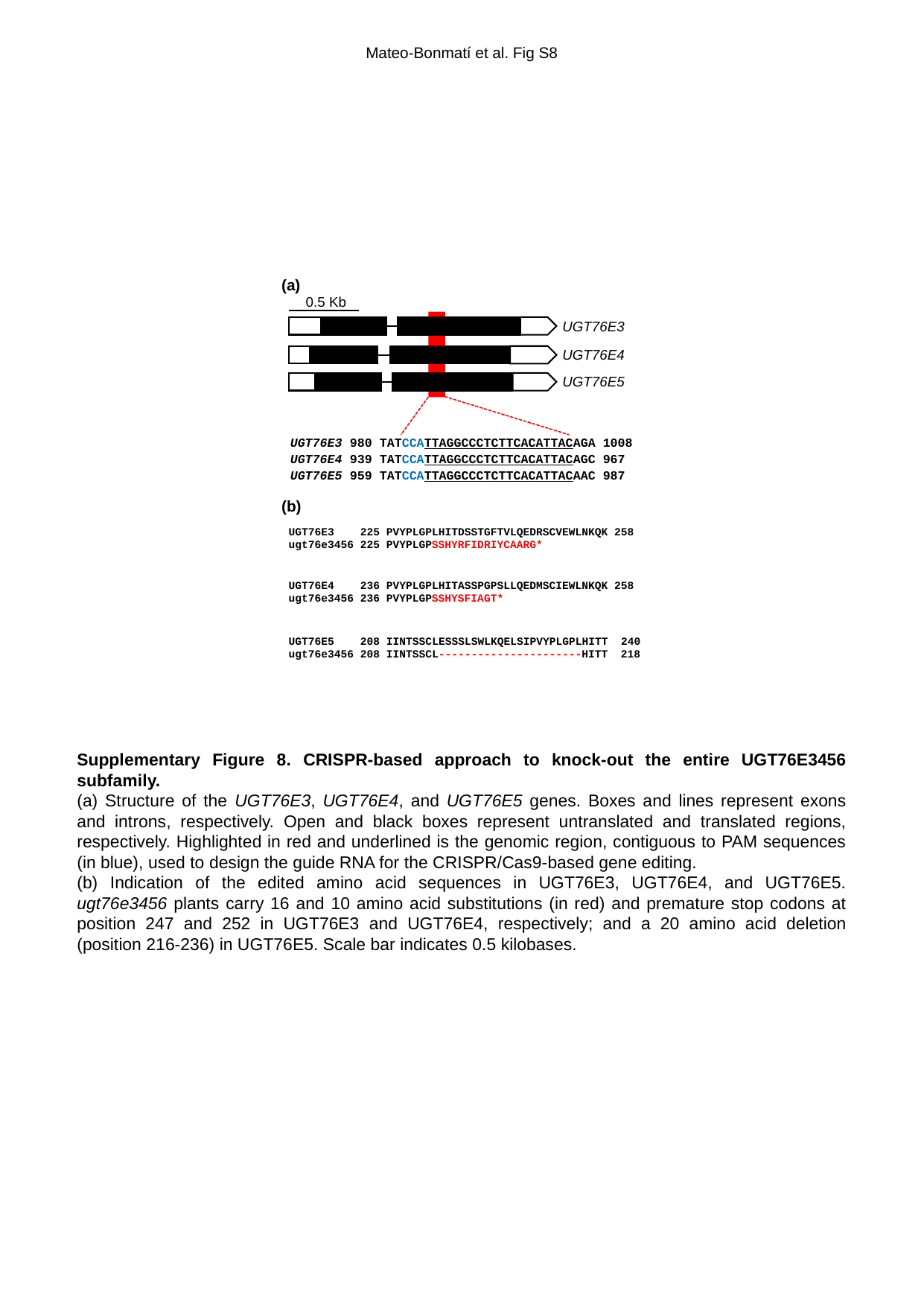

Mateo-Bonmatí et al. Fig S8
(a)
0.5 Kb
UGT76E3
UGT76E4
UGT76E5
UGT76E3 980 TATCCATTAGGCCCTCTTCACATTACAGA 1008
UGT76E4 939 TATCCATTAGGCCCTCTTCACATTACAGC 967
UGT76E5 959 TATCCATTAGGCCCTCTTCACATTACAAC 987
(b)
UGT76E3 225 PVYPLGPLHITDSSTGFTVLQEDRSCVEWLNKQK 258
ugt76e3456 225 PVYPLGPSSHYRFIDRIYCAARG*
UGT76E4 236 PVYPLGPLHITASSPGPSLLQEDMSCIEWLNKQK 258
ugt76e3456 236 PVYPLGPSSHYSFIAGT*
UGT76E5 208 IINTSSCLESSSLSWLKQELSIPVYPLGPLHITT 240
ugt76e3456 208 IINTSSCL----------------------HITT 218
Supplementary Figure 8. CRISPR-based approach to knock-out the entire UGT76E3456 subfamily.
(a) Structure of the UGT76E3, UGT76E4, and UGT76E5 genes. Boxes and lines represent exons and introns, respectively. Open and black boxes represent untranslated and translated regions, respectively. Highlighted in red and underlined is the genomic region, contiguous to PAM sequences (in blue), used to design the guide RNA for the CRISPR/Cas9-based gene editing.
(b) Indication of the edited amino acid sequences in UGT76E3, UGT76E4, and UGT76E5. ugt76e3456 plants carry 16 and 10 amino acid substitutions (in red) and premature stop codons at position 247 and 252 in UGT76E3 and UGT76E4, respectively; and a 20 amino acid deletion (position 216-236) in UGT76E5. Scale bar indicates 0.5 kilobases.

### Slide 11
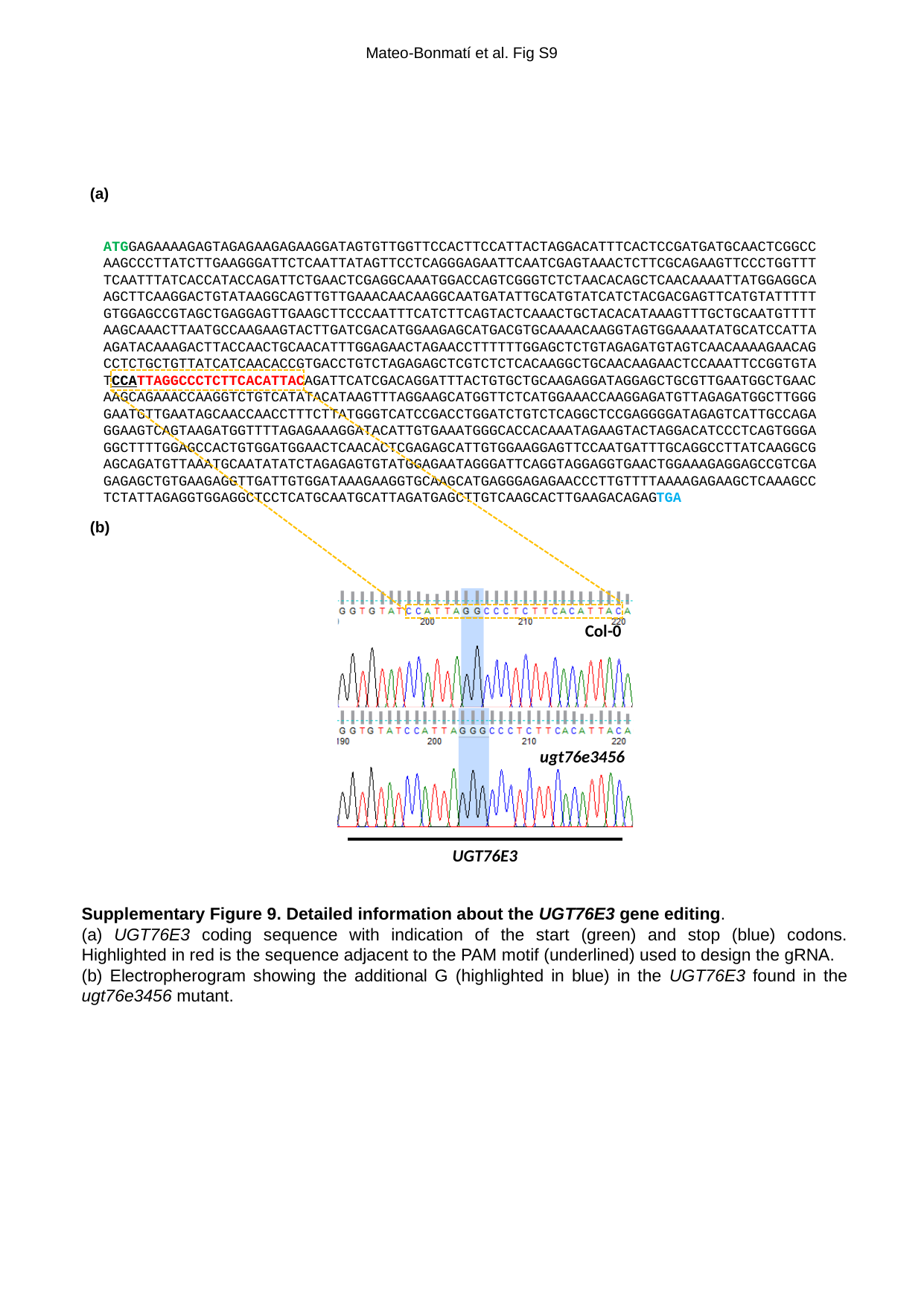

Mateo-Bonmatí et al. Fig S9
(a)
ATGGAGAAAAGAGTAGAGAAGAGAAGGATAGTGTTGGTTCCACTTCCATTACTAGGACATTTCACTCCGATGATGCAACTCGGCCAAGCCCTTATCTTGAAGGGATTCTCAATTATAGTTCCTCAGGGAGAATTCAATCGAGTAAACTCTTCGCAGAAGTTCCCTGGTTTTCAATTTATCACCATACCAGATTCTGAACTCGAGGCAAATGGACCAGTCGGGTCTCTAACACAGCTCAACAAAATTATGGAGGCAAGCTTCAAGGACTGTATAAGGCAGTTGTTGAAACAACAAGGCAATGATATTGCATGTATCATCTACGACGAGTTCATGTATTTTTGTGGAGCCGTAGCTGAGGAGTTGAAGCTTCCCAATTTCATCTTCAGTACTCAAACTGCTACACATAAAGTTTGCTGCAATGTTTTAAGCAAACTTAATGCCAAGAAGTACTTGATCGACATGGAAGAGCATGACGTGCAAAACAAGGTAGTGGAAAATATGCATCCATTAAGATACAAAGACTTACCAACTGCAACATTTGGAGAACTAGAACCTTTTTTGGAGCTCTGTAGAGATGTAGTCAACAAAAGAACAGCCTCTGCTGTTATCATCAACACCGTGACCTGTCTAGAGAGCTCGTCTCTCACAAGGCTGCAACAAGAACTCCAAATTCCGGTGTATCCATTAGGCCCTCTTCACATTACAGATTCATCGACAGGATTTACTGTGCTGCAAGAGGATAGGAGCTGCGTTGAATGGCTGAACAAGCAGAAACCAAGGTCTGTCATATACATAAGTTTAGGAAGCATGGTTCTCATGGAAACCAAGGAGATGTTAGAGATGGCTTGGGGAATGTTGAATAGCAACCAACCTTTCTTATGGGTCATCCGACCTGGATCTGTCTCAGGCTCCGAGGGGATAGAGTCATTGCCAGAGGAAGTCAGTAAGATGGTTTTAGAGAAAGGATACATTGTGAAATGGGCACCACAAATAGAAGTACTAGGACATCCCTCAGTGGGAGGCTTTTGGAGCCACTGTGGATGGAACTCAACACTCGAGAGCATTGTGGAAGGAGTTCCAATGATTTGCAGGCCTTATCAAGGCGAGCAGATGTTAAATGCAATATATCTAGAGAGTGTATGGAGAATAGGGATTCAGGTAGGAGGTGAACTGGAAAGAGGAGCCGTCGAGAGAGCTGTGAAGAGGTTGATTGTGGATAAAGAAGGTGCAAGCATGAGGGAGAGAACCCTTGTTTTAAAAGAGAAGCTCAAAGCCTCTATTAGAGGTGGAGGCTCCTCATGCAATGCATTAGATGAGCTTGTCAAGCACTTGAAGACAGAGTGA
(b)
Col-0
ugt76e3456
UGT76E3
Supplementary Figure 9. Detailed information about the UGT76E3 gene editing.
(a) UGT76E3 coding sequence with indication of the start (green) and stop (blue) codons. Highlighted in red is the sequence adjacent to the PAM motif (underlined) used to design the gRNA.
(b) Electropherogram showing the additional G (highlighted in blue) in the UGT76E3 found in the ugt76e3456 mutant.

### Slide 12
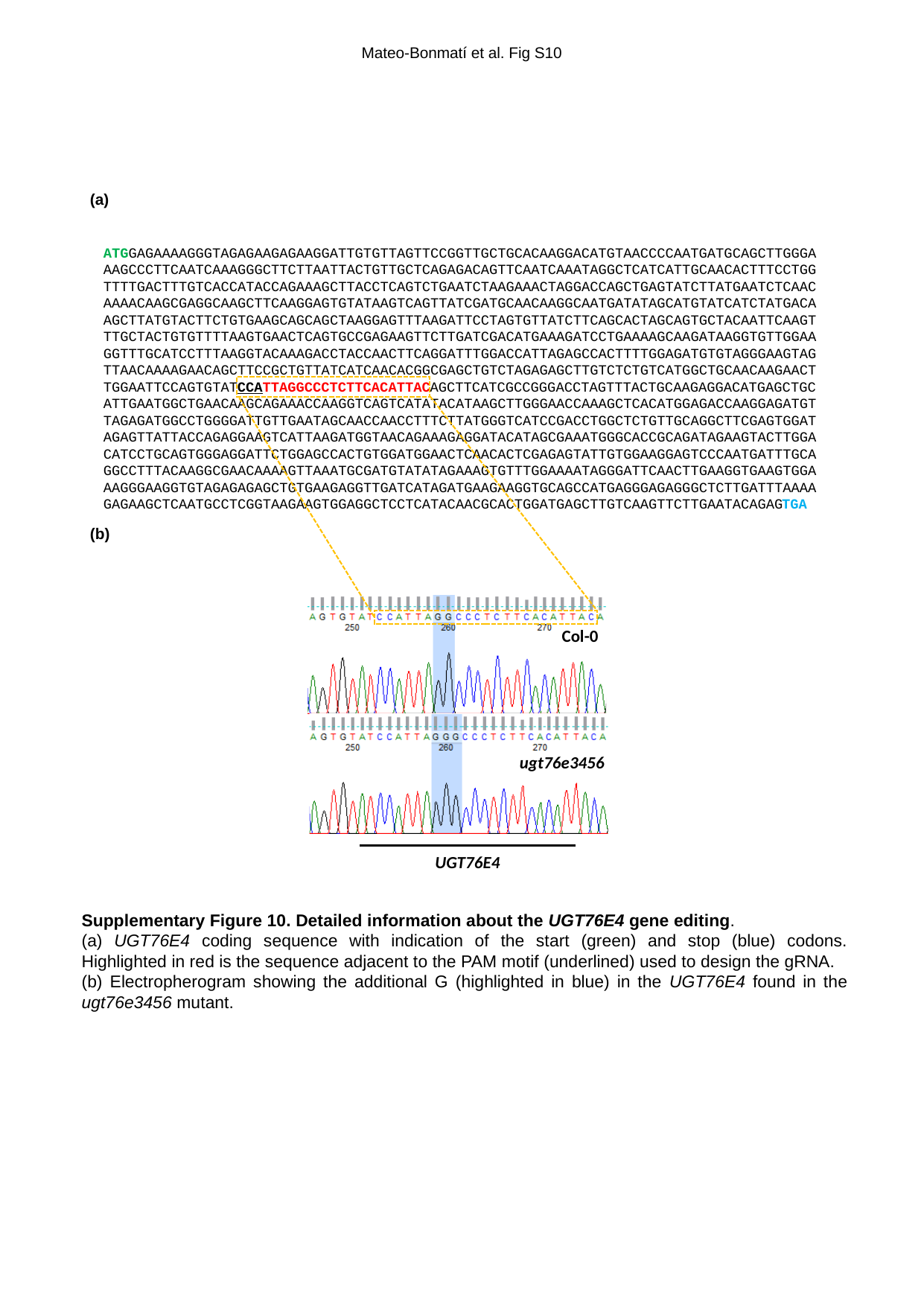

Mateo-Bonmatí et al. Fig S10
(a)
ATGGAGAAAAGGGTAGAGAAGAGAAGGATTGTGTTAGTTCCGGTTGCTGCACAAGGACATGTAACCCCAATGATGCAGCTTGGGAAAGCCCTTCAATCAAAGGGCTTCTTAATTACTGTTGCTCAGAGACAGTTCAATCAAATAGGCTCATCATTGCAACACTTTCCTGGTTTTGACTTTGTCACCATACCAGAAAGCTTACCTCAGTCTGAATCTAAGAAACTAGGACCAGCTGAGTATCTTATGAATCTCAACAAAACAAGCGAGGCAAGCTTCAAGGAGTGTATAAGTCAGTTATCGATGCAACAAGGCAATGATATAGCATGTATCATCTATGACAAGCTTATGTACTTCTGTGAAGCAGCAGCTAAGGAGTTTAAGATTCCTAGTGTTATCTTCAGCACTAGCAGTGCTACAATTCAAGTTTGCTACTGTGTTTTAAGTGAACTCAGTGCCGAGAAGTTCTTGATCGACATGAAAGATCCTGAAAAGCAAGATAAGGTGTTGGAAGGTTTGCATCCTTTAAGGTACAAAGACCTACCAACTTCAGGATTTGGACCATTAGAGCCACTTTTGGAGATGTGTAGGGAAGTAGTTAACAAAAGAACAGCTTCCGCTGTTATCATCAACACGGCGAGCTGTCTAGAGAGCTTGTCTCTGTCATGGCTGCAACAAGAACTTGGAATTCCAGTGTATCCATTAGGCCCTCTTCACATTACAGCTTCATCGCCGGGACCTAGTTTACTGCAAGAGGACATGAGCTGCATTGAATGGCTGAACAAGCAGAAACCAAGGTCAGTCATATACATAAGCTTGGGAACCAAAGCTCACATGGAGACCAAGGAGATGTTAGAGATGGCCTGGGGATTGTTGAATAGCAACCAACCTTTCTTATGGGTCATCCGACCTGGCTCTGTTGCAGGCTTCGAGTGGATAGAGTTATTACCAGAGGAAGTCATTAAGATGGTAACAGAAAGAGGATACATAGCGAAATGGGCACCGCAGATAGAAGTACTTGGACATCCTGCAGTGGGAGGATTCTGGAGCCACTGTGGATGGAACTCAACACTCGAGAGTATTGTGGAAGGAGTCCCAATGATTTGCAGGCCTTTACAAGGCGAACAAAAGTTAAATGCGATGTATATAGAAAGTGTTTGGAAAATAGGGATTCAACTTGAAGGTGAAGTGGAAAGGGAAGGTGTAGAGAGAGCTGTGAAGAGGTTGATCATAGATGAAGAAGGTGCAGCCATGAGGGAGAGGGCTCTTGATTTAAAAGAGAAGCTCAATGCCTCGGTAAGAAGTGGAGGCTCCTCATACAACGCACTGGATGAGCTTGTCAAGTTCTTGAATACAGAGTGA
(b)
Col-0
ugt76e3456
UGT76E4
Supplementary Figure 10. Detailed information about the UGT76E4 gene editing.
(a) UGT76E4 coding sequence with indication of the start (green) and stop (blue) codons. Highlighted in red is the sequence adjacent to the PAM motif (underlined) used to design the gRNA.
(b) Electropherogram showing the additional G (highlighted in blue) in the UGT76E4 found in the ugt76e3456 mutant.

### Slide 13
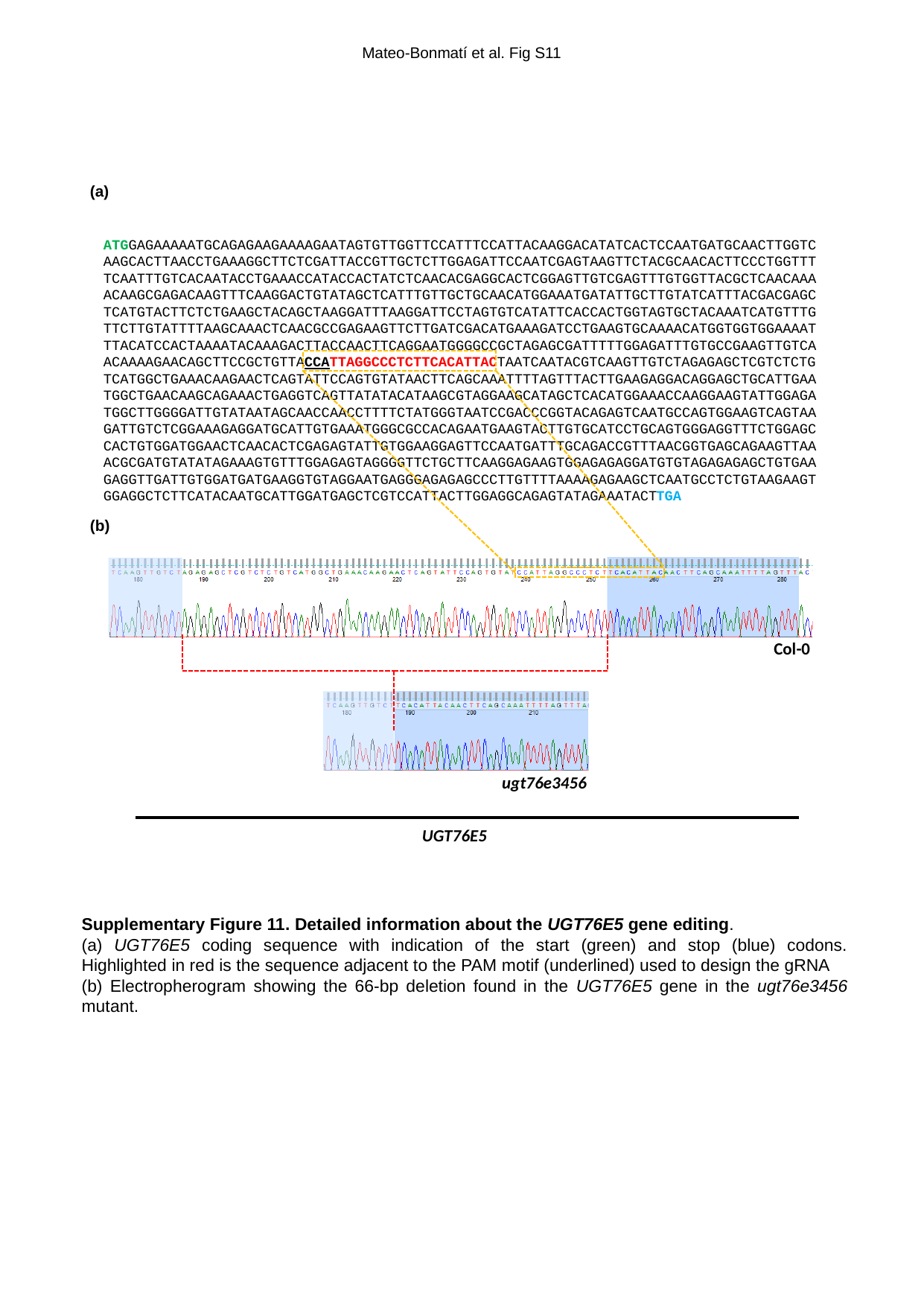

Mateo-Bonmatí et al. Fig S11
(a)
ATGGAGAAAAATGCAGAGAAGAAAAGAATAGTGTTGGTTCCATTTCCATTACAAGGACATATCACTCCAATGATGCAACTTGGTCAAGCACTTAACCTGAAAGGCTTCTCGATTACCGTTGCTCTTGGAGATTCCAATCGAGTAAGTTCTACGCAACACTTCCCTGGTTTTCAATTTGTCACAATACCTGAAACCATACCACTATCTCAACACGAGGCACTCGGAGTTGTCGAGTTTGTGGTTACGCTCAACAAAACAAGCGAGACAAGTTTCAAGGACTGTATAGCTCATTTGTTGCTGCAACATGGAAATGATATTGCTTGTATCATTTACGACGAGCTCATGTACTTCTCTGAAGCTACAGCTAAGGATTTAAGGATTCCTAGTGTCATATTCACCACTGGTAGTGCTACAAATCATGTTTGTTCTTGTATTTTAAGCAAACTCAACGCCGAGAAGTTCTTGATCGACATGAAAGATCCTGAAGTGCAAAACATGGTGGTGGAAAATTTACATCCACTAAAATACAAAGACTTACCAACTTCAGGAATGGGGCCGCTAGAGCGATTTTTGGAGATTTGTGCCGAAGTTGTCAACAAAAGAACAGCTTCCGCTGTTACCATTAGGCCCTCTTCACATTACTAATCAATACGTCAAGTTGTCTAGAGAGCTCGTCTCTGTCATGGCTGAAACAAGAACTCAGTATTCCAGTGTATAACTTCAGCAAATTTTAGTTTACTTGAAGAGGACAGGAGCTGCATTGAATGGCTGAACAAGCAGAAACTGAGGTCAGTTATATACATAAGCGTAGGAAGCATAGCTCACATGGAAACCAAGGAAGTATTGGAGATGGCTTGGGGATTGTATAATAGCAACCAACCTTTTCTATGGGTAATCCGACCCGGTACAGAGTCAATGCCAGTGGAAGTCAGTAAGATTGTCTCGGAAAGAGGATGCATTGTGAAATGGGCGCCACAGAATGAAGTACTTGTGCATCCTGCAGTGGGAGGTTTCTGGAGCCACTGTGGATGGAACTCAACACTCGAGAGTATTGTGGAAGGAGTTCCAATGATTTGCAGACCGTTTAACGGTGAGCAGAAGTTAAACGCGATGTATATAGAAAGTGTTTGGAGAGTAGGGGTTCTGCTTCAAGGAGAAGTGGAGAGAGGATGTGTAGAGAGAGCTGTGAAGAGGTTGATTGTGGATGATGAAGGTGTAGGAATGAGGGAGAGAGCCCTTGTTTTAAAAGAGAAGCTCAATGCCTCTGTAAGAAGTGGAGGCTCTTCATACAATGCATTGGATGAGCTCGTCCATTACTTGGAGGCAGAGTATAGAAATACTTGA
(b)
Col-0
ugt76e3456
UGT76E5
Supplementary Figure 11. Detailed information about the UGT76E5 gene editing.
(a) UGT76E5 coding sequence with indication of the start (green) and stop (blue) codons. Highlighted in red is the sequence adjacent to the PAM motif (underlined) used to design the gRNA
(b) Electropherogram showing the 66-bp deletion found in the UGT76E5 gene in the ugt76e3456 mutant.

### Slide 14
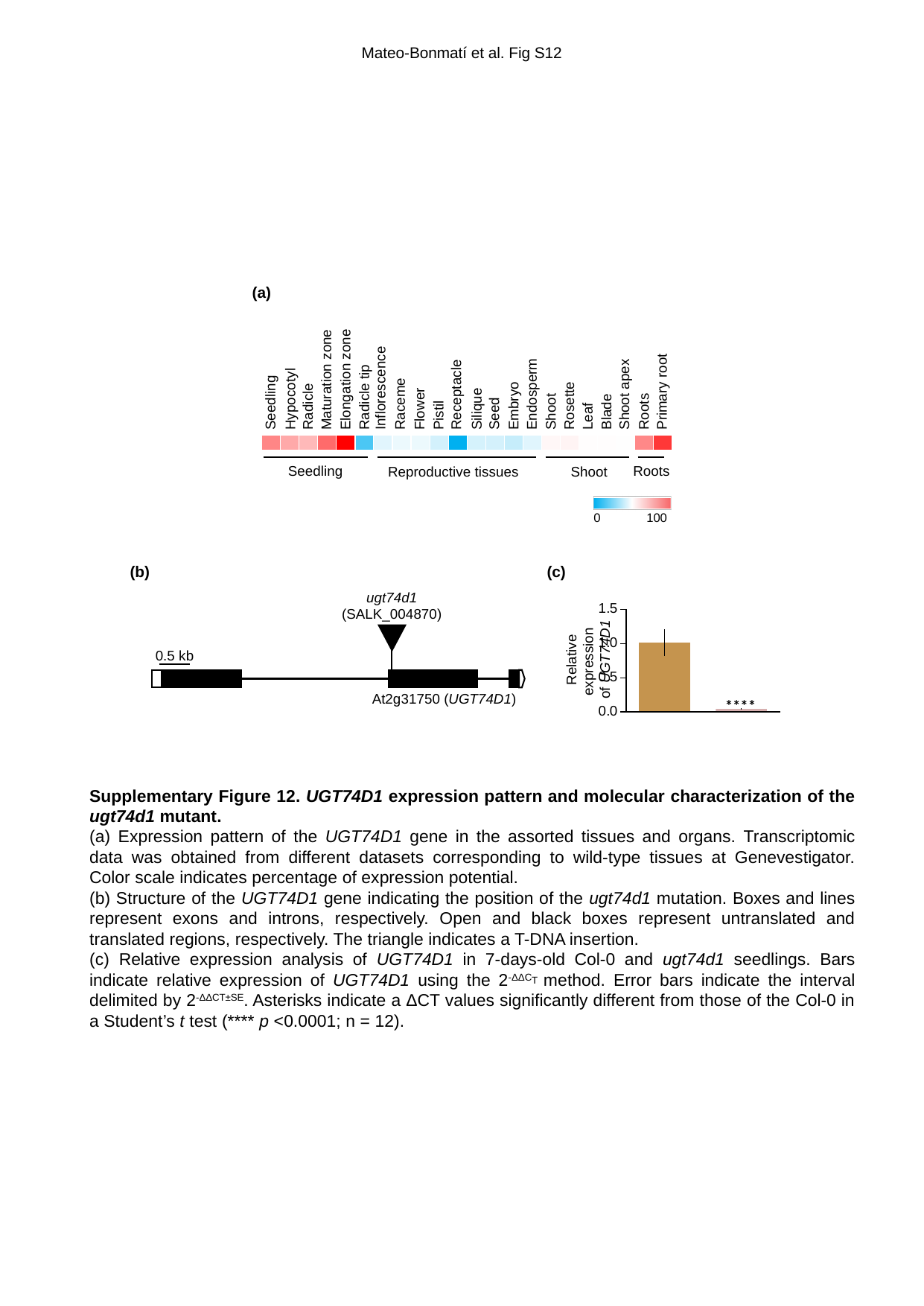

Mateo-Bonmatí et al. Fig S12
(a)
Maturation zone
Elongation zone
Inflorescence
Embryo
Endosperm
Shoot
Rosette
Leaf
Blade
Shoot apex
Roots
Primary root
Receptacle
Radicle tip
Hypocotyl
Seedling
Raceme
Radicle
Silique
Flower
Seed
Pistil
Seedling
Roots
Reproductive tissues
Shoot
0
100
(b)
(c)
ugt74d1
(SALK_004870)
Relative expression
of UGT74D1
0.5 kb
At2g31750 (UGT74D1)
****
Supplementary Figure 12. UGT74D1 expression pattern and molecular characterization of the ugt74d1 mutant.
(a) Expression pattern of the UGT74D1 gene in the assorted tissues and organs. Transcriptomic data was obtained from different datasets corresponding to wild-type tissues at Genevestigator. Color scale indicates percentage of expression potential.
(b) Structure of the UGT74D1 gene indicating the position of the ugt74d1 mutation. Boxes and lines represent exons and introns, respectively. Open and black boxes represent untranslated and translated regions, respectively. The triangle indicates a T-DNA insertion.
(c) Relative expression analysis of UGT74D1 in 7-days-old Col-0 and ugt74d1 seedlings. Bars indicate relative expression of UGT74D1 using the 2^-ΔΔCT^ method. Error bars indicate the interval delimited by 2^-ΔΔCT±SE^. Asterisks indicate a ΔCT values significantly different from those of the Col-0 in a Student’s t test (**** p <0.0001; n = 12).

### Slide 15
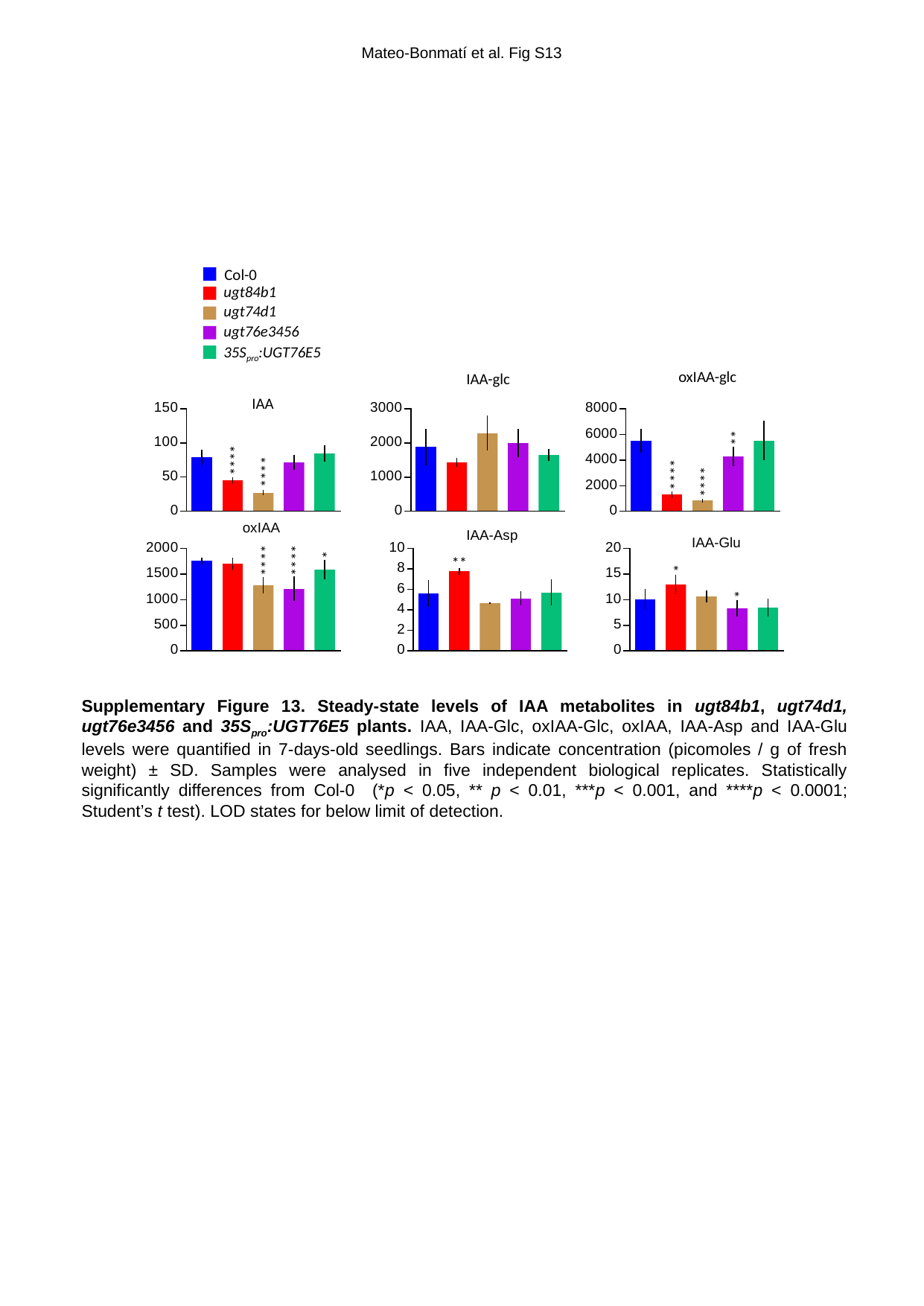

Mateo-Bonmatí et al. Fig S13
Col-0
ugt84b1
ugt74d1
ugt76e3456
35S_pro_:UGT76E5
oxIAA-glc
IAA-glc
IAA
**
****
****
****
****
oxIAA
IAA-Asp
IAA-Glu
*
****
****
**
*
*
Supplementary Figure 13. Steady-state levels of IAA metabolites in ugt84b1, ugt74d1, ugt76e3456 and 35S_pro_:UGT76E5 plants. IAA, IAA-Glc, oxIAA-Glc, oxIAA, IAA-Asp and IAA-Glu levels were quantified in 7-days-old seedlings. Bars indicate concentration (picomoles / g of fresh weight) ± SD. Samples were analysed in five independent biological replicates. Statistically significantly differences from Col-0 (*p < 0.05, ** p < 0.01, ***p < 0.001, and ****p < 0.0001; Student’s t test). LOD states for below limit of detection.

### Slide 16
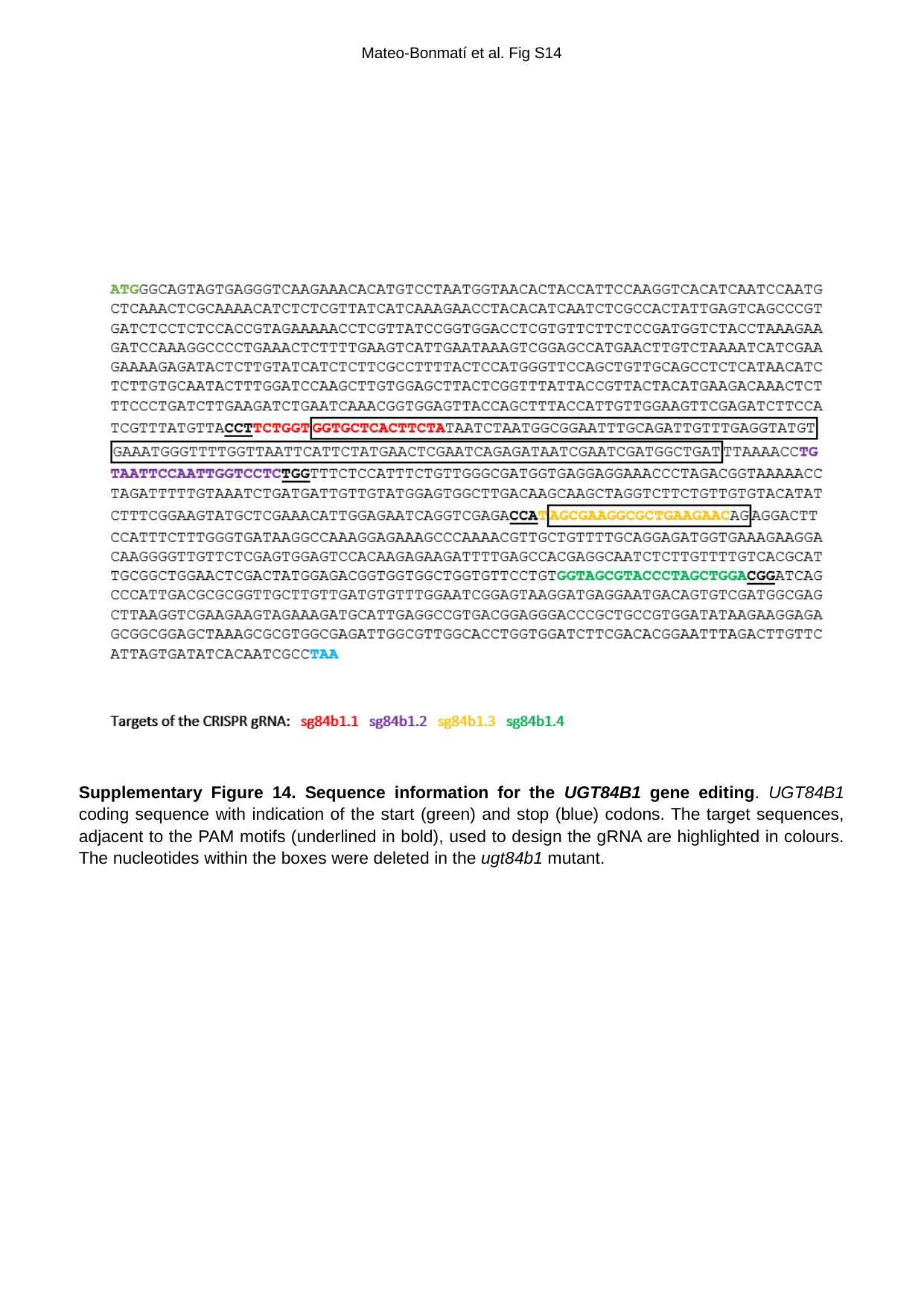

Mateo-Bonmatí et al. Fig S14
Supplementary Figure 14. Sequence information for the UGT84B1 gene editing. UGT84B1 coding sequence with indication of the start (green) and stop (blue) codons. The target sequences, adjacent to the PAM motifs (underlined in bold), used to design the gRNA are highlighted in colours. The nucleotides within the boxes were deleted in the ugt84b1 mutant.
